# Predicting Cell Movement for Low-Frame-Rate Tracking

**DOI:** 10.64898/2026.11.20.768128

**Authors:** Mina Gachloo, Tirthankar Biswas, Bogumila Ndubuisi, Marc R. Birtwistle, Federico Iuricich

## Abstract

Automated cell tracking is widely used to quantify cellular behavior and typically involves two steps: identifying cells in each observation and linking corresponding cells across observations. This paper focuses on the linking problem under low-frame-rate imaging. Low-frame-rate imaging mitigates photobleaching and phototoxicity and enables long-term observations, but larger cell displacements make spatial proximity less reliable and increase ambiguity among candidate links. We investigate whether cell images contain information about future movement direction that can provide an additional cue for cell linking. We systematically evaluate the impact of image-patch configurations, direction representations, and deep-learning architectures for movement-direction prediction accuracy. Moreover, we incorporate the predicted directions into a Linear Assignment Problem (LAP)-based linking method and evaluate linking performance across four cell lines and multiple temporal gaps. Our results show that movement-direction information can improve cell linking by 2.7 to 5.7 percentage points over the distance-only LAP baseline, depending on the cell line, at a two-hour interval between frames. Among the evaluated configurations, a CNN–LSTM with a class-based direction representation provides the largest improvement (4.1 percentage points on average) over the distance-only LAP baseline. These results show that cell images provide a cue for cell linking that spatial proximity alone does not capture.

## 1 Introduction

Automated analysis of individual cells in time-lapse microscopy supports quantitative studies of cell migration and morphological dynamics [16, 49], as well as cell proliferation and lineage reconstruction [32]. Time-lapse imaging is also used to study intracellular dynamics and cellular responses to experimental perturbations [25, 31]. These measurements support a wide range of applications, including drug-response screening [10, 43], wound-healing and cell-migration assays [23], and studies of stem-cell differentiation and reprogramming [8, 38].

However, microscopy experiments face an important trade-off between temporal resolution and experimental quality. Frequent image acquisition provides detailed information about cell movement and cellular events, but repeated illumination increases phototoxicity [27] and photobleaching [42]. High acquisition rates can also limit the number of fields of view or experimental conditions that can be measured simultaneously, reducing the scalability of high-throughput experiments [45].

Reducing the acquisition frequency can mitigate these limitations and enable longer or larger-scale experiments. However, sparse acquisition creates a more difficult cell-linking problem [17, 18]. Cells may move considerable distances, change their appearance, or undergo division between consecutive observations. As a result, spatial proximity becomes less reliable, and multiple candidate cells in the next frame appear equally plausible. This motivates our central question: can information extracted from cell images provide a complementary cue that improves linking beyond a distance-based LAP baseline?

In this work, we investigate whether deep-learning models can infer a cell’s subsequent movement direction from image patches and whether incorporating these predictions into the linking process improves cell linking under sparse acquisition. Rather than proposing a new linking algorithm, we present a systematic methodological study of the design choices involved in movement prediction, including the input provided to the model, the representation used for the predicted direction, and the architecture used to process the image information. Our contributions are:

- a controlled perturbation analysis that quantifies how movement-direction errors affect cell-linking recall as a function of the temporal gap;
- a systematic comparison of four image-patch input configurations that combine the current, past, and future patches for movement prediction;
- an evaluation of discrete direction classification and continuous angular regression as alternative prediction formulations;
- an assessment of multiple deep-learning architectures for processing and combining image information; and
- an evaluation of the best-performing configurations within a linear-assignment-based linking method across progressively larger temporal gaps.

## 2 Related Work

This section first reviews methods for computing cell links, establishing the problem addressed in this work (Section 2.1). We then discuss approaches for predicting cell movement, which can provide additional information for resolving ambiguous links (Section 2.2). Finally, we describe the challenges introduced by low-frame-rate imaging and review methods developed to address them (Section 2.3).

### 2.1 Computing cell links

Many cell-tracking methods follow a tracking-by-detection framework, in which tracking is separated into two stages: detecting cells in individual frames and linking corresponding cells across frames [34, 36, 51]. The linking step is often formulated as a Linear Assignment Problem (LAP) [24, 48], where a cost function quantifies the dissimilarity between candidate cell pairs. This cost may be based on spatial relationships, such as the distance between cell positions [14, 24], or on the overlap between detected cell regions [5]. When spatial information alone is insufficient, methods may additionally compare appearance features such as intensity or cell size [24, 48]. These approaches require dataset-specific design choices: users must determine how spatial and appearance information are combined, assign weights to the different cues, and select thresholds for excluding measurements considered unreliable [19].

Deep learning methods reduce the need to manually design appearance features and tune their relative weights. Instead, these representations are learned from annotated data [34, 36, 37].

Some approaches work directly on raw images and integrate both cell identification and cell linking. A recent example is CellTrack R-CNN which analyzes pairs of adjacent frames using a Mask R-CNN-based detection and segmentation network [12]. It then compares candidate cells across frames using both their visual appearance and relative spatial arrangement, producing a similarity score that indicates whether two detections correspond to the same cell.

Cell-TRACTR also performs detection, segmentation, and linking within a single model [40]. It first locates and segments cells in an image. For subsequent frames, the learned representation associated with each previously detected cell is converted into a track query and passed to a transformer together with the new image. The transformer uses this query to locate the corresponding cell and predict its position and segmentation mask in the new frame.

CELLECT follows a similar strategy for 3D microscopy but is designed for positional annotations rather than dense segmentation masks [53]. Given adjacent image frames, it predicts candidate cell centers and learns a feature representation for each center. Candidate centers in neighboring frames are then compared using their learned representations together with spatial information to determine whether they represent the same cell, a division event, or different cells.

In contrast, other approaches assume that cell instances have already been detected in each frame. They learn a cost or association score for each candidate cell pair and pass these scores to a separate optimization procedure, such as the LAP, which determines the final links [19, 37].

An early example is Caliban, which employs a neural network to predict, for each pair of cells in consecutive frames, the probability that they are the same cell, different cells, or parent and daughter, and uses these probabilities as costs in the LAP [37].

Ben-Haim and Riklin Raviv instead formulate linking as a graph-based classification problem: cells are represented as nodes, candidate links as edges, and a graph neural network predicts which edges correspond to valid cell associations [2].

More recently, Trackastra uses a transformer to predict associations among cells within a local temporal window [19]. Each detection is represented by its position and shallow features computed from the corresponding image region or segmentation, including mean intensity, area, and shape descriptors. These features are combined with learned spatial and temporal encodings and provided to the transformer, which predicts an association probability for each pair of detections. Predictions from overlapping temporal windows are averaged and used to construct a candidate graph. Trackastra can then obtain the final cell links through an optimization-based linker (e.g., LAP).

### 2.2 Motion prediction

A complementary strategy for improving cell linking is to model cell motion and use it to estimate each cell’s likely future position.

Traditional approaches describe cell movement using predefined motion models. For example, under a Brownian-motion model [1,14], the displacement distribution over a given time interval defines a high-probability region around the cell’s current position. This region can be used to constrain linking: detections outside it are assigned a low likelihood under the model and excluded as candidate links.

A Kalman filter [26] extends this idea by using observations from previous frames to maintain and update a cell-specific estimate of motion. At each time point, the filter propagates the previous state (i.e., cell position and velocity) through the motion model to predict the cell’s position in the next frame. When a new detection becomes available, the prediction is updated by combining the predicted state with the observation while accounting for the uncertainty associated with each. The predicted position and its uncertainty then define a search region for cell linking: detections within this region are retained as candidate links, whereas detections outside it are assigned a low likelihood and excluded [24, 48, 50]. Motion estimates can also incorporate contextual information, such as the local cell density or the number of cells within a neighborhood, to adapt the predicted movement to the surrounding population [35].

Beyond predefined motion models, some methods infer cell movement directly from image appearance using deep learning. Nishimoto et al. [39] showed that a CNN can predict the subsequent direction of cell motion from a single image patch by assigning it to one of four directional quadrants. The method achieved accuracies between 82% and 95%, depending on the cell line. Visualization of the network’s predictions highlighted morphological cues associated with migration, including protrusions and the trailing edge. Because four quadrants provide only a coarse description of movement direction, Bruns et al. [9] formulated direction estimation as a continuous angular regression problem. Their model predicts an angle on a circular domain, where 0*^◦^* and 360*^◦^* denote the same direction. This formulation achieved a mean angular error of approximately 17*^◦^* on two datasets, compared with 30*^◦^*and 34*^◦^*, respectively, for the quadrant-based baseline evaluated on the same datasets.

Motion prediction can also be integrated directly into cell identification and linking. Hayashida et al. introduced the Motion and Position Map (MPM), which jointly predicts a map indicating likely cell locations and a motion vector field that points toward the corresponding cell center in the neighboring frame [21]. The predicted vectors therefore provide both evidence that a cell is present and a direct correspondence between cells across frames. Their subsequent method incorporates multiple frames and introduces an object-level warping loss, which penalizes motion predictions that are inconsistent when propagated across time [22].

EmbedTrack follows a related strategy using a CNN that processes two consecutive frames. In addition to predicting the cell regions in each frame, the network predicts, for every cell pixel in the current frame, an offset toward the center of the corresponding cell in the previous frame. These offsets are used to identify the corresponding instances and establish the links between them [30]. DiSTNet2D extends motion-based linking to a larger temporal context by processing seven temporally distributed frames and predicting forward and backward displacement information for the cells [41]. The resulting motion estimates are used during post-processing to establish links and to correct segmentation errors that are inconsistent with the observed temporal behavior.

Taken together, these studies demonstrate the value of motion information for cell linking. However, most have been developed and evaluated on sequences with relatively short acquisition intervals and limited displacement between consecutive observations. For example, many commonly used Cell Tracking Challenge datasets have acquisition intervals Δ*t* of 5 to 30 minutes [51], This leaves open how effectively motion-based information can support cell linking when the inter-frame displacements is larger.

### 2.3 Low-frame-rate cell tracking

Linking methods and motion models generally perform reliably when image sequences are densely sampled [34, 47, 51]. Under sparse acquisition, however, cells can move substantially between consecutive observations, while divisions, cell entry or exit, and other events may occur between acquisitions without being observed. These effects increase the ambiguity of cell associations across frames [17, 54]. Consequently, linking performance typically decreases as the interval between observations increases, with the severity of this degradation depending on factors such as cell motility and population density [17, 18, 54].

These challenges have motivated approaches that either reconstruct the missing temporal information or exploit structural information that remains available across sparsely sampled frames. Priessner et al. [44] use deep-learning-based frame interpolation to generate intermediate images between acquired frames, enabling computational recovery of temporal information that was not recorded. Their method is evaluated on microscopy sequences with acquisition intervals ranging from seconds to minutes.

Chen et al. [13] instead use the spatial organization of cells to establish associations without relying on an explicit motion model. Their method constructs a Delaunay triangulation of the cell positions in each frame and links cells by matching corresponding Delaunay subgraphs across frames.

Alternatively, other methods incorporate motion information to account for larger displacements. Hayashida and Bise [20] estimate a Cell Motion Field from consecutive images, which encodes the motion of each cell along the segment joining its centroids in the two frames. This estimated motion is combined with the distance between candidate cells to determine their links. The method is evaluated on low-frame-rate sequences with a 25-minute interval between consecutive frames.

More recently, Bragantini et al. [7] incorporate temporal registration into Ultrack to estimate local motion between frames. Nonlinear registration produces a movement vector field that is applied to candidate segmentations during tracking. The approach is evaluated on the Cell Tracking Challenge Tribolium dataset [34] at temporal intervals of 1.5, 7.5, and 15 minutes, respectively.

The method proposed by Zyss et al. [54] learns representations that capture cell appearance and morphology. A contrastive model brings observations of the same cell closer in representation space while separating observations of different cells. During linking, similarities between these representations are combined with spatial and additional appearance-based features to define edges in a graph, from which cell tracks are recovered through graph optimization. Although the method reports strong performance on sequences acquired at intervals of up to four hours, some of the evaluated sequences exhibit relatively small inter-frame displacements despite their long acquisition intervals. In addition, the implementation is not publicly available [54].

## 3 Methods

This section describes the experimental framework used to systematically investigate deep-learning-based movement predictions as an additional source of information for cell linking in low-frame-rate microscopy.

Our central premise is that predicting a cell’s future movement can provide valuable information for resolving cell links and improving cell-linking accuracy. Specifically, we investigate whether cell images contain cues that predict subsequent movement and whether incorporating these predictions into the cell-linking process improves the accuracy of cell links.

Our analysis is organized around four components: the input, the output representation, and the network architecture of movement-direction prediction, and the effect of the prediction on cell linking. First, we evaluate different image-patch configurations to determine which spatial and temporal context should be provided as input (Figure 1a, Section 3.1). Second, we compare two output formulations, a class-based representation, where the direction is predicted as one of K discrete classes,and a continuous-angle representation, where the direction is predicted as a continuous angle (Figure 1b, Section 3.2). Third, we assess how different network architectures process and combine the available image information (Figure 1c, Section 3.3). Finally, we evaluate the effect of the predicted direction on cell linking by incorporating the predicted movement into a linking algorithm (Figure 1d, Section 3.4).

**Figure 1:**
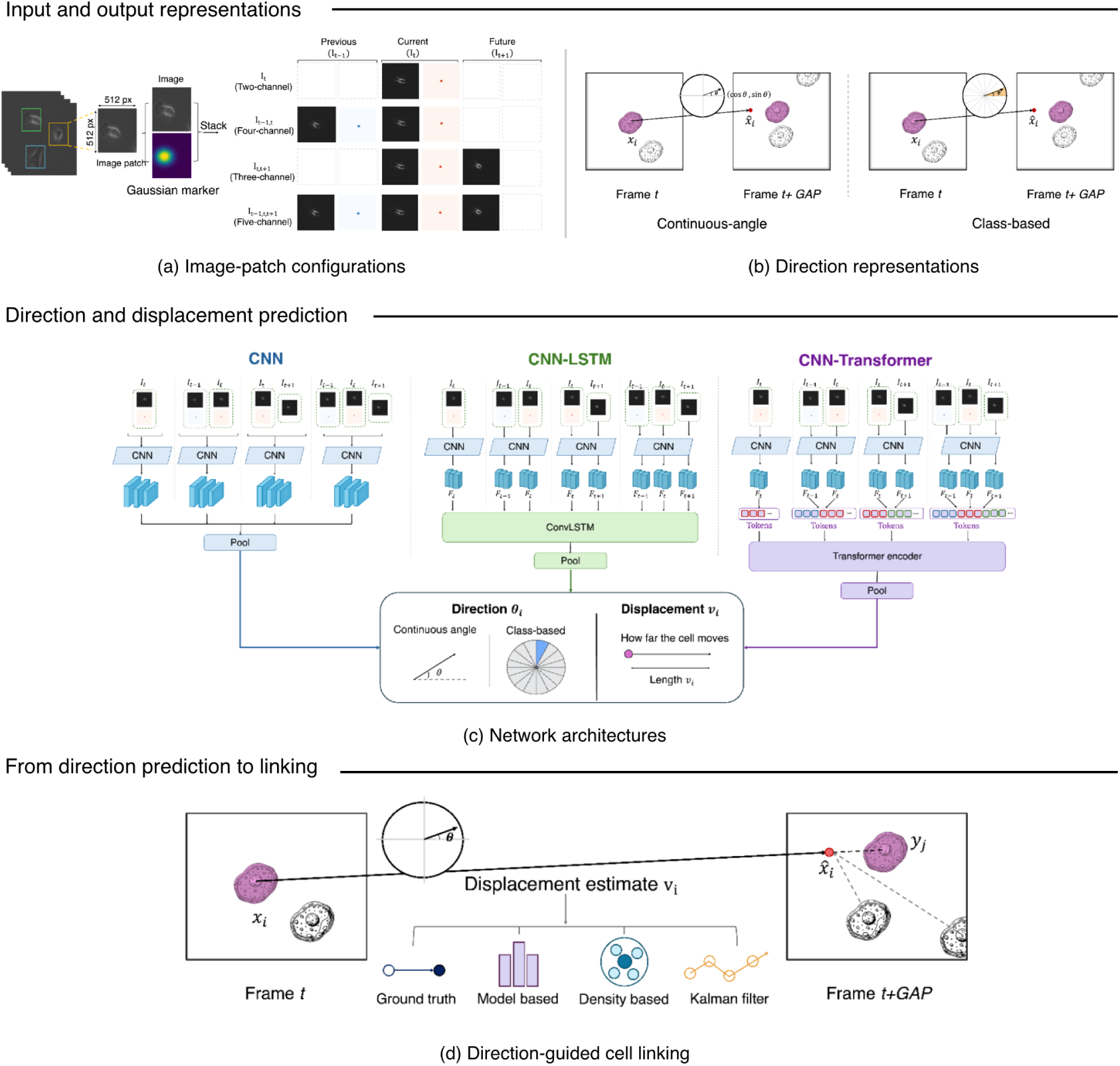
Overview of our methodology for investigating image-based movement predictions to improve cell linking. The study examines three design choices. (a) **Image-patch configurations.** Different types of image patches are extracted around each cell from consecutive frames, with Gaussian marker channels indicating the target cell. (b) **Direction representations.** Movement direction is represented either as a continuous angle or as a discrete direction class. (c) **Network architectures.** A CNN, a CNN–LSTM, and a CNN–Transformer are used to predict movement direction and displacement from the image patches. Finally, (d) the predicted direction is used to project each cell to its expected position, using one of four displacement estimates (ground truth, model based, density based, and Kalman filter), and the resulting positions are used to compute linking costs.

### 3.1 Image-Patch Configurations

Our first question is what spatial and temporal information should be provided to the model to predict a cell’s future movement. We assume a cell-linking framework in which the cell’s current position and its previously established links are available. Based on this information, we consider three possible sources of image context:

- the current patch, extracted from the source frame;
- a past patch, extracted from the previous frame;
- and a future patch, extracted from the future frame.

All three patches are extracted with the same 512 × 512 window, centered on the cell’s current position, from the phase-contrast images of the LFCT dataset (Section 4.1). The future patch uses this position because the cell’s future position is unknown before linking, and the past patch uses it so that the cell’s displacement from the previous frame remains visible. This fixed window lets the model compare cell appearance and position within the same image region across frames. The window size is chosen to account for the increased cell displacement at larger temporal gaps.

Based on these three patch types, we construct four input configurations that combine the available spatial and temporal information in different ways. These configurations let us evaluate the contribution of each patch type individually and in combination.

Since a patch may contain several cells, the current and past patches are paired with a second channel containing a Gaussian marker that identifies the target cell. In the current patch, the marker is centered on the cell’s current position. In the past patch, the marker indicates the cell’s previously observed position, which is available from the established link. No marker is provided for the future patch because the cell’s future position is unknown.

The collection of patches provided to the model defines the four configurations evaluated in this work (Figure 1a):

- *I_t_* - contains the patch extracted at the current frame and its marker. The movement direction is inferred from the appearance of the cell at a single time point.
- *I_t−_*_1*,t*_ - contains the patches extracted at the current frame and the previous frame, adding the appearance of the cell before the current frame.
- *I_t,t_*_+1_ - contains the patches extracted at the current frame and the future frame, adding the region of the future frame into which the cell has moved.
- *I_t−_*_1*,t,t*+1_ - contains all three patches, providing past, current, and future information.

### 3.2 Movement Direction Representation

Our second question is how to represent movement direction to better support the learning process. Previous work has explored the idea of predicting motion directions either as a discrete class [39] or as a continuous angle [9]. However, these two formulations have not been directly compared under a common experimental setting. Our focus is to compare the class-based and continuous angular regression within the same linking framework with particular attention to their performance at larger temporal gaps. Figure 1b illustrates the two representations.

#### Discrete class representation

In this setting, the movement direction output is represented as a discrete label. The circular direction space is divided into *N* equal classes, each covering 360*^◦^/N* . Assigning an angle to its nearest class center introduces a maximum quantization error of half the class width. In our experimental evaluation, we use *N* = {4, 8, 16, 32, 64}, corresponding to class widths of 90*^◦^*, 45*^◦^*, 22.5*^◦^*, 11.25*^◦^*, and 5.625*^◦^*, respectively. For each choice of *N* , the class-based network (see Section 3.3) is adapted by setting its final fully connected layer to *N* output units, with one logit corresponding to each direction class. A softmax converts these logits into class probabilities. The class with the highest probability is selected, and the center angle of its corresponding angular sector is used as the predicted movement direction.

#### Continuous angle representation

In this setting, the movement direction output is represented as a continuous angle. Directly regressing the angle is affected by its periodicity because directions close to 0*^◦^* and 360*^◦^*are identical but numerically distant. To handle this periodicity, the network predicts the two-dimensional unit vector (cos *θ,* sin *θ*) representing the direction angle *θ* [4, 9]. In the regression network this is implemented by a final fully connected layer containing two units which output is normalized to unit length. The predicted angle is recovered from the two components using the two-argument arctangent function. The model is trained using a cosine distance loss between the predicted and ground-truth unit vectors, which is minimized when the predicted and true directions coincide [4].

### 3.3 Network Architectures for Direction Prediction

Our third question is how different network architectures perform for direction prediction. We evaluate three architectures that process the frames of a configuration in different ways: a CNN stacks them as input channels, a CNN-LSTM processes them sequentially through recurrence, and a CNN-Transformer relates them through attention. In all three, the resulting features are passed through fully connected layers that produce the direction representation described in Section 3.2. Each architecture also includes a separate output head that predicts the displacement *v_i_*. The three architectures are illustrated in Figure 1c.

The first architecture is a CNN [28, 29], in which all frames and markers are stacked as input channels and processed together in a single pass. The network begins with a 7 × 7 convolution with 32 filters, followed by max pooling. This is followed by three convolutional blocks with 64, 128, and 256 filters, respectively. Each block contains two 3 × 3 convolutions, with the first using a stride of 2 to reduce the spatial resolution. Normalization and ReLU activation are applied after each convolution. The resulting feature map is reduced using adaptive average pooling.

The second architecture is a CNN-LSTM based on a convolutional LSTM (ConvLSTM) [46], in which the frames are processed sequentially through recurrence. Each frame input is first processed by a shared ResNet-18 encoder pretrained on ImageNet, with its first convolution adapted to two input channels. The encoder is truncated after the third residual block, producing a spatial feature map with 256 channels for each frame. These feature maps are projected to 128 channels using a 1×1 convolution and passed sequentially through a ConvLSTM with 128 hidden channels and a 3 × 3 convolutional kernel. The ConvLSTM maintains hidden and cell states as spatial feature maps, preserving spatial information as the sequence is processed. The final hidden state is passed through a 3 × 3 convolution followed by adaptive average pooling.

The third architecture is a CNN-Transformer [52], in which the frames are related through attention. Each frame input is first processed by a shared ResNet-18 encoder pretrained on ImageNet, with its first convolution adapted to two input channels. The encoder is truncated after the fourth residual block, producing a spatial feature map with 512 channels for each frame. These feature maps are projected to 128 channels using a 1 × 1 convolution, and each spatial location is treated as a token. Spatial and temporal positional encodings are added to the tokens to preserve their location within each frame and their order across frames. The tokens are then processed by four Transformer blocks with four attention heads. Each block applies divided space-time attention [3], in which temporal attention first relates tokens at the same spatial location across frames, followed by spatial attention among the tokens within each frame. After the attention blocks, the tokens corresponding to the current frame are reshaped into a spatial feature map and passed through a 3 × 3 convolution followed by adaptive average pooling.

### 3.4 From direction prediction to linking

To go beyond direction prediction accuracy and explicitly estimate the impact on linking accuracy we use a Linear Assignment Problem (LAP)-based technique for linking. Not only this approach is well-established and easy to implement [24, 48] but it also demonstrated the most robust performance when temporal gaps are introduced [17] (Figure 4b).

To avoid confounding factors such as mitosis, cell entry or exit, and apoptosis, we restrict the LAP to the known sets of cells present in the two frames being linked. That is, we only link cells that are present in both frames, so that every cell in frame *t* has its correct match in frame *t* + 1.

Given the sets of cells *X* = {*x_i_*} in frame *t* and *Y* = {*y_j_*} in frame *t* + 1, LAP works on a cost matrix

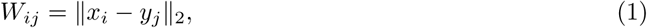

where ∥ · ∥ _2_ denotes the Euclidean distance between cell positions. The Hungarian algorithm solves this assignment problem, producing the assignment that minimizes the total cost, with each source cell paired to at most one target cell.

To incorporate movement predictions into the assignment problem, we replace each cell’s current position with its predicted position in the future frame. For a cell *x_i_* in frame *t*, let *θ_i_* and *v_i_* denote the predicted movement direction and displacement, respectively. The predicted position is

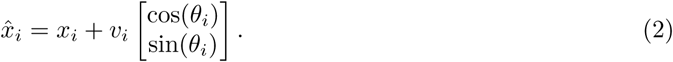

The angle prediction *θ_i_* is computed by the networks discussed in Section 3.3. The velocity *v* is estimated following four different approaches:

- *Ground truth* - The model uses the ground-truth displacement magnitude for each cell. Although this quantity is unavailable in practical applications, this oracle configuration provides a way to measures the improvement in cell linking that could be achieved if the displacement were known exactly.
- *Model based* - Both the movement direction *θ_i_* and displacement *v_i_* are predicted by the CNN-LSTM using input *I_t−_*_1*,t,t*+1_. The displacement is predicted by a separate output head of the network (Section 3.3). This architecture and input configuration are selected based on the experiments presented in Section 4.3.1. The two predictions are combined using Equation 2.
- *Density based* - The displacement *v_i_*is estimated according to the number *n_i_* of other cells within a fixed radius of cell *i*. Cells are assigned to three groups: few (*n_i_* ≤ 3), medium (*τ*_1_ *< n_i_* ≤ 7), and many (*n_i_ >* 7). For each density group, we first estimate the distribution of displacement magnitudes from the training links. We then select a single representative value from the corresponding displacement range, choosing the value that minimizes linking error on the training sequences. At test time, a cell assigned to density group *g* is given the corresponding value *v_g_*, which is then used as its predicted displacement.
- *Kalman filter* - The displacement is estimated from the links the tracker has already produced. Each cell carries an estimate *v*^ of its displacement together with an uncertainty. When a link is formed with observed displacement, the estimate is updated using a one-dimensional Kalman filter [26],

For each of these approaches, once *θ_i_* and *v_i_* are defined for all cells in *X*, the cost of assigning a source cell *x_i_* to a target cell *y_j_*is computed as the Euclidean distance between the predicted and target positions:

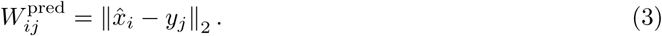

## 4 Experimental evaluation

In this Section we discuss the results obtained in our experiments. We first present our experimental framework in Section 4.1, including the dataset, the training procedure, and the evaluation metrics. Then, in Section 4.2, we study how errors in the predicted direction affect cell linking under controlled conditions. Finally, in Section 4.3, we present the cell linking results obtained with the movement predictions of our models.

The results in this section focus on GAP 8, the highest sparsification level considered in this work, corresponding to a two-hour acquisition interval between consecutive frames. Results for smaller gaps are provided in the supplementary material.

### 4.1 Experimental framework

#### Dataset

All experiments are conducted on the Low-Frame-Rate Cell Tracking (LFCT) dataset [17], which contains multi-day time-lapse microscopy sequences of four human cell lines. The dataset provides a total of 25 sequences across four cell lines: 9 MCF10A, 6 HEK293T, 6 MDA-MB-231, and 4 U87. We use image sequences acquired at 20× magnification where each cell occupies more pixels, so the morphological cues associated with migration are better resolved, and where relative inter-frame movement for each cell is larger [17]. These sequences are sparsified to simulate low frame rate using GAP factors 1, 2, 4, and 8, where GAP *x* retains every *x*-th frame of the original sequence. Frames are acquired every 15 minutes, so the interval between retained frames increases from 15 minutes at GAP 1 to 2 hours at GAP 8. Figure 2 shows the inter-frame displacement distributions for the 20× sequences. The median displacement increases with the temporal gap for all four cell lines. These larger displacements make spatial proximity less reliable for identifying corresponding cells across frames. The complete dataset and ground-truth annotations are available at https://davislab.github.io/LFCT-dataset/.

**Figure 2:**
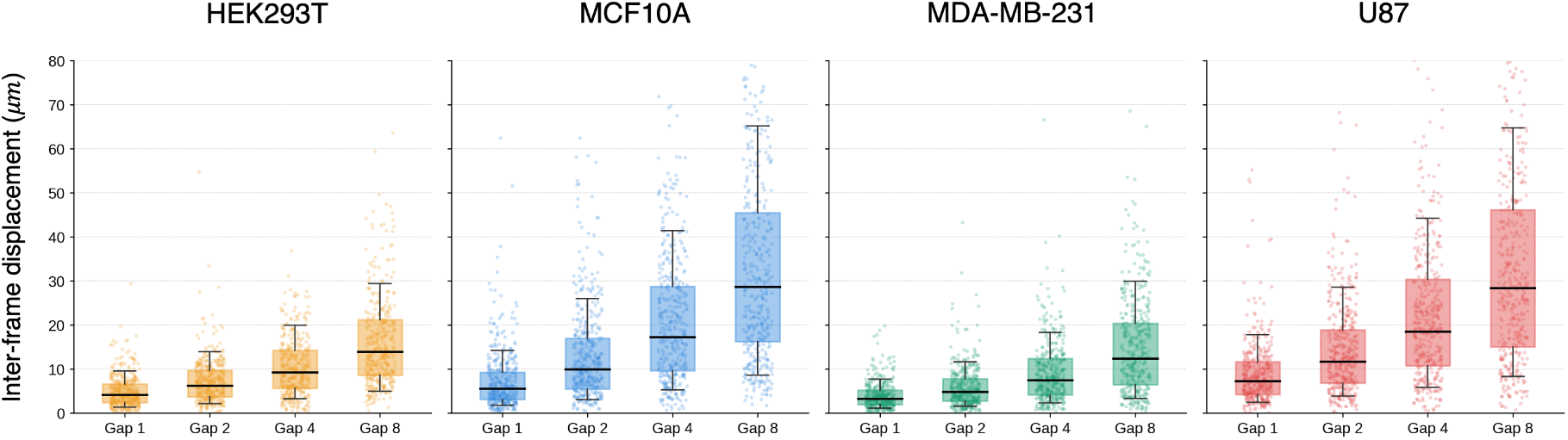
Inter-frame displacement across temporal gaps for the 20× sequences of the LFCT dataset, with one panel per cell line. Boxes show the interquartile range, black lines indicate the median, and whiskers show the 10th and 90th percentiles. Dots show sampled displacement observations.

#### Training

All models are trained for 50 epochs with the AdamW optimizer and evaluated with three-fold cross-validation. Models are trained separately for each cell line. Within a cell line, one sequence is held out for validation and three sequences rotate through the test position, with the remaining sequences used for training. Folds are defined by image sequence, so within each fold no sequence appears in more than one split, and the checkpoint is selected on the validation sequence.

Hyperparameters are tuned separately for each architecture, and their values are provided in the configuration file of the code repository.

#### Evaluation metrics

Results are analyzed using two metrics: angular error and linking accuracy. We use *angular error* for both discrete and continuous predictions. For continuous angular regression, this is the circular difference between the predicted angle *θ*^^^ and the ground-truth angle *θ*:

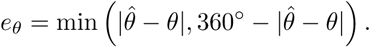

For discrete predictions, we convert the predicted class into its corresponding representative angle before computing the same circular error. This metric accounts for the periodicity of angular values, such that predictions near 0*^◦^*and 360*^◦^* are treated as close. We primarily compare the distributions of angular errors across models rather than relying only on their average values.

Finally, we evaluate the effect of predicted movement on cell linking in a fixed assignment setting (see Section 3.4, Figure 1d). Ground-truth links are taken from the manually curated annotations of the LFCT dataset [17]. The evaluation considers only links in which a cell in frame *t* is connected to the same cell in the subsequent frame *t* + 1; division links are excluded. A predicted link is counted as a true positive when it connects the same source and target cells as the ground-truth correspondence. A ground-truth link that is not recovered is counted as a false negative. Accuracy is evaluated by *linking recall*, which measures the fraction of true links that are correctly recovered:

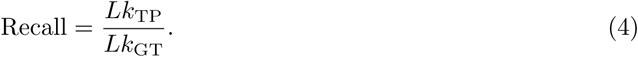

where *Lk*_TP_ is the number of correctly recovered links and *Lk*_GT_ is the total number of ground-truth links.

### 4.2 Controlled analysis of direction-prediction errors

Before evaluating learned movement predictions, we first ask how accurate a predicted direction must be to improve cell linking over the distance-only LAP baseline. To answer this, we start from the ground-truth movement directions, introduce progressively larger amounts of angular noise, and use the perturbed directions to predict cell positions for linking.

The resulting relationship between prediction error and linking performance indicates the level of directional accuracy required for movement predictions to improve cell linking, and is used in the following sections to interpret the errors of the learned models.

Perturbed directions are generated starting from the ground-truth direction *θ* by sampling circular noise from a von Mises distribution [15, 33]. The perturbed direction is *θ̃* = wrap_[0,2*π*)_ (*θ* + *ɛ_σ_*) where:

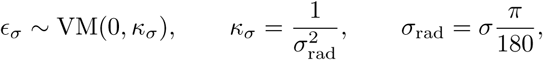

and *σ* denotes the noise spread in degrees.

Each perturbation is an independent random draw, and errors larger than *σ* may occur. We evaluate *σ* ∈ {0, 10, 20, 30, 45, 60, 90, 120, 135, 150, 180}*^◦^*, repeating each noise level 20 times with independent draws and averaging the resulting linking performance.

Figure 3a shows the distributions of the angular noise generated by the von Mises noise model for selected values of *σ*. At small *σ* the noise concentrates near 0*^◦^*, while at large *σ* the distribution becomes nearly uniform.

**Figure 3:**
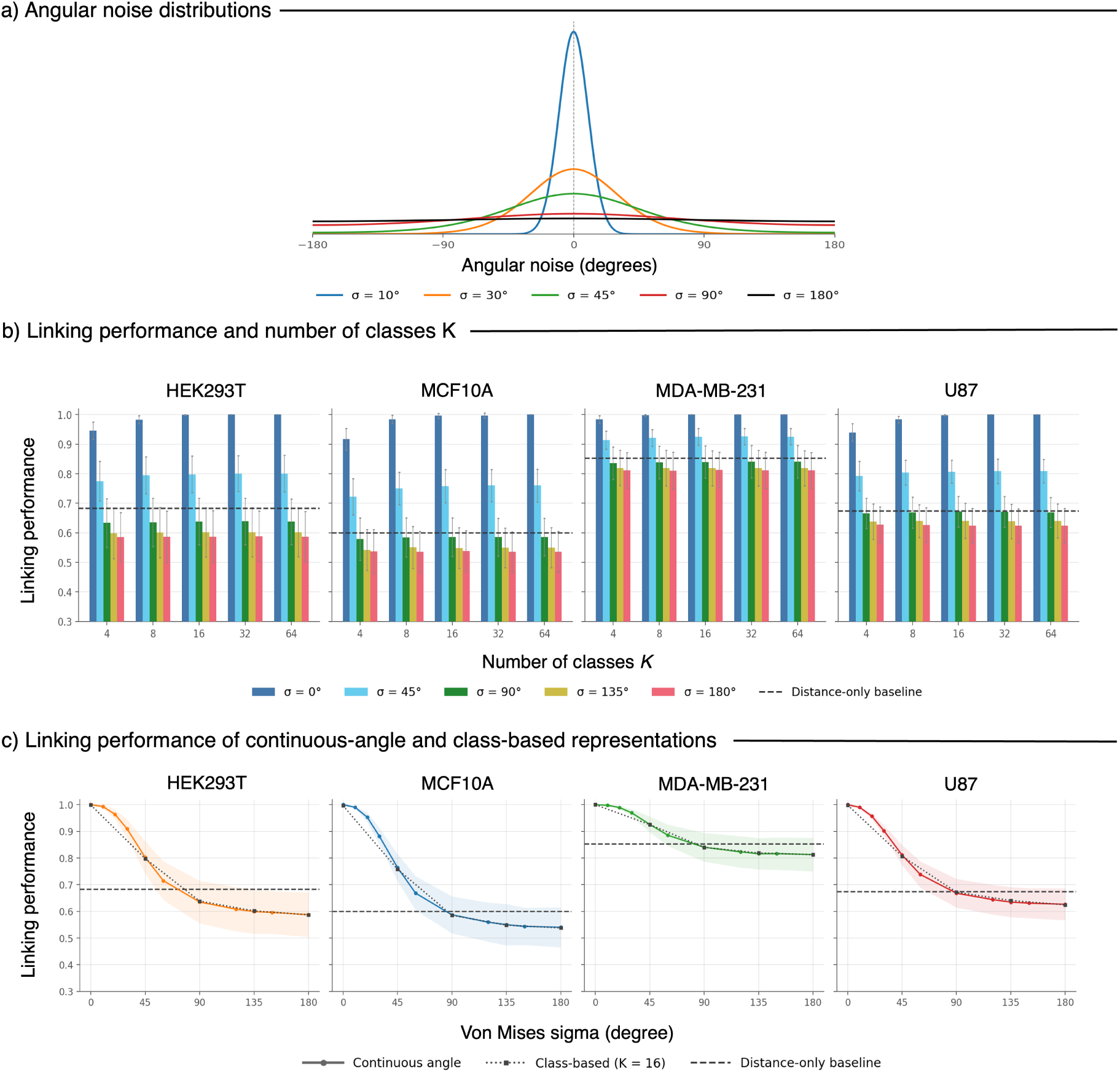
Controlled analysis of direction-prediction errors at GAP 8. (a) Distribution of the angular error *θ̃* − *θ* introduced by the von Mises noise model for selected values of *σ*, where *θ* is the ground-truth direction and *θ̃* the perturbed direction. (b) Linking performance when the perturbed directions are quantized into *K* classes, one panel per cell line. Bars show the mean across fields of view and error bars the standard deviation. (c) Linking performance for the continuous-angle and class-based (*K* = 16) representations as a function of *σ*. Lines show the mean across fields of view and shaded bands the standard deviation. In (b) and (c), the dashed line shows the distance-only LAP baseline.

**Figure 4:**
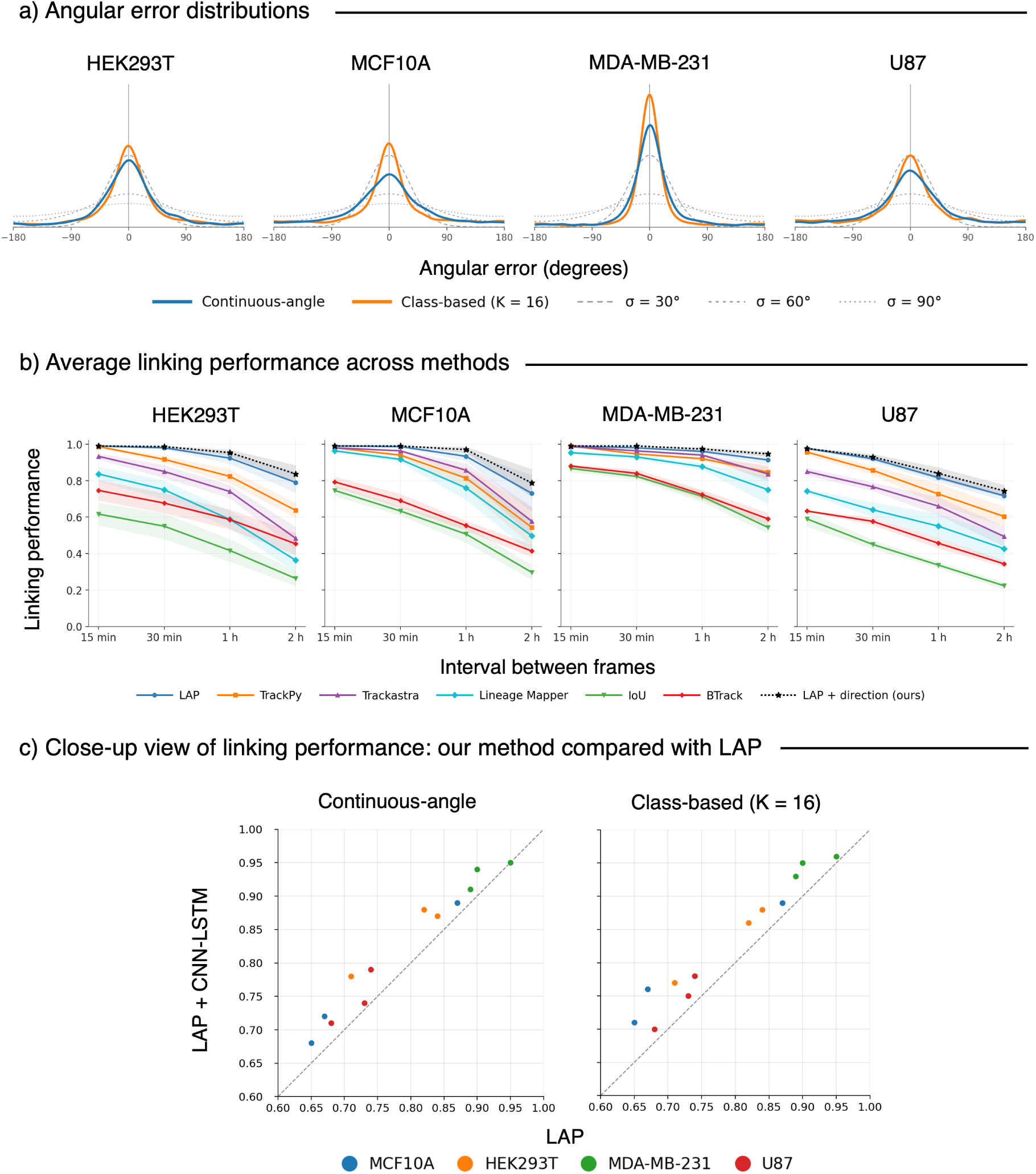
(a) Angular error distributions of the CNN–LSTM with input *I_t−_*_1*,t,t*+1_ at GAP 8 for the continuous-angle and class-based (*K* = 16) representations, with von Mises reference distributions at *σ* = 30*^◦^*, 60*^◦^*, and 90*^◦^*. (b) Linking performance of six cell-linking methods across temporal gaps (15 min to 2 h) for each cell line: the LAP, TrackPy [1], Trackastra [19], Lineage Mapper [11], the intersection-over-union tracker (IoU) [6], and the Bayesian Tracker (BTrack) [50]. The black dotted line shows our approach, LAP with movement directions predicted by the CNN-LSTM using the class-based representation. The lines show the average across sequences of the same cell line. Shaded regions represent variability across sequences of the same cell line. (c) Linking performance of the configurations in (a) at GAP 8 with the density based displacement, compared with the distance-only LAP baseline. Each point is one test sequence, and points above the diagonal indicate an improvement.

To relate direction-prediction error to linking performance, we compute cell links using the LAP-based approach. We use the perturbed direction for *θ* and the ground-truth displacement magnitude for *v*, ensuring that any reduction in linking performance can be attributed solely to errors in the predicted direction.

For the continuous angular formulation, *θ̃* is used directly in the predicted-position equation. For the class-based formulation with *K* direction classes, the perturbed angle is converted to its corresponding direction class. The direction domain is divided into *K* equal sectors of width Δ = 2*π/K*, and *θ̃* is assigned to the sector containing it, *k* = ⌊*θ̃ /*Δ⌋, where *k* ∈ {0*, . . . , K* − 1}. The representative angle of the assigned sector, *θ_k_* = *k* + ^1^ Δ, is then used in the predicted-position equation.

Before comparing discrete and continuous direction prediction, we first characterize how the number of direction classes affects linking performance. For several noise levels (*σ*), we quantize the perturbed ground-truth directions into *K* ∈ {4, 8, 16, 32, 64} uniformly spaced classes and use the corresponding class representatives to compute the LAP assignments.

#### Optimal number of classes *K*

Figure 3b shows linking performance as a function of the number of direction classes for the evaluated noise levels. The largest improvement with increasing resolution occurs from *K* = 4 to *K* = 8, particularly at low noise levels. With four classes, each class spans 90*^◦^*, and much of the directional information is lost through this discretization alone. A smaller improvement remains from *K* = 8 to *K* = 16 at *σ* = 0*^◦^*, while increasing the resolution to *K* = 32 or *K* = 64 brings little further change. Based on these results we select *K* = 16 for all subsequent experiments.

#### Linking performance in controlled scenarios

Figure 3c compares linking performance across noise levels for the continuous-angle and class-based direction representations. The dashed line indicates the baseline performance obtained by the distance-only LAP method. We observe no substantial difference in linking performance between the continuous-angle and class-based representations across the evaluated noise levels. This suggests that, in principle, both formulations can provide comparable linking performance when they produce predictions of similar accuracy. Their practical difference will then depend on how robustly a given model can learn each type of information from image data. For both representations, linking performance is generally below the distance-only LAP baseline at noise levels of *σ* ≥ 90*^◦^*, whereas it typically exceeds the baseline when *σ* ≤ 60*^◦^*. Thus, a movement-direction predictor with an error distribution narrower than that produced by the *σ* = 60*^◦^*perturbation should generally be expected to improve linking performance over the distance-only baseline.

### 4.3 Cell Linking with movement predictions

In this section we present the complete results of our analysis. We start by showcasing the cell linking results of the best performing model among those we have explored compared to the LAP baseline (see Section 3.4). Then we proceed discussing the relative improvement due to image patches, model architecture, and cell velocity estimation.

#### 4.3.1 Best performing models

In this section we present cell linking results using the best-performing model and input configurations identified in our experiments. A detailed analysis of each configuration choice is discussed in the following sections.

For both direction representations, the best performing model was the CNN–LSTM architecture using image-patch configuration *I_t−_*_1*,t,t*+1_ as input. For the class-based representation, we use *K* = 16 direction classes. The density based configuration performed best for estimating displacement.

Figure 4a shows the angular-error distributions of these configurations for both direction representations, together with reference distributions at selected noise levels from the controlled analysis. For all cell lines, the class-based representation produces predictions with a lower median angular error (9.2*^◦^* to 23.5*^◦^*) compared to the continuous angle representation (13.9*^◦^* to 32.8*^◦^*). All these values are below the median angular error of about 49*^◦^* produced by the *σ* = 60*^◦^* perturbation, the noise level up to which linking performance typically exceeds the distance-only LAP baseline (Section 4.2).

Figure 4b shows the linking performance of six linking approaches as a function of the temporal gap for four different cell lines. Performance decreases as the gap increases for all methods, but the size of the decrease varies across methods and cell lines. Interestingly, a simple LAP-based linking method is the most robust approach to increasing gaps. Adding the predicted movement direction to LAP further improves linking performance, and the improvement is largest at the longest gap.

We next evaluate how these direction errors affect cell linking. Using the density based displacement, both representations improve linking performance compared with the distance-only LAP baseline (Figure 4c). On average, linking performance increases by 3.4 percentage points with the continuous-angle representation and by 4.1 points with the class-based representation. This pattern is consistent with the lower angular errors of the class-based representation. For the class-based representation, the average improvement per cell line ranges from 2.7 points for U87 to 5.7 points for MCF10A.

#### 4.3.2 Predicting direction from image patches

The first contribution to the improvement seen in Section 4.3.1 comes from the input patch configuration. Beyond the single-patch input already used in the literature [39], past and future patches improve the ability of the network to predict movement direction.

Figure 5a shows the error distribution for each input configuration of the CNN–LSTM at GAP 8. With a single image *I_t_*, the distribution is almost flat for most cell lines, with a median angular error between 55*^◦^* and 82*^◦^*for the continuous-angle models and between 33*^◦^* and 73*^◦^*for the class-based models, indicating that one image is not enough to predict movement over an interval of two hours. Adding past information *I_t−_*_1*,t*_ reduces the median error only slightly for most cell lines. When future frames are involved, the predictions concentrate close to the true direction: the median angular error decreases to 14–36*^◦^* for the continuous-angle models and to 9–25*^◦^* for the class-based models with *I_t,t_*_+1_ and *I_t−_*_1*,t,t*+1_, and the fraction of predictions within 22.5*^◦^*of the true direction increases from 15–38% with *I_t_* to 38–76% with *I_t−_*_1*,t,t*+1_.

**Figure 5:**
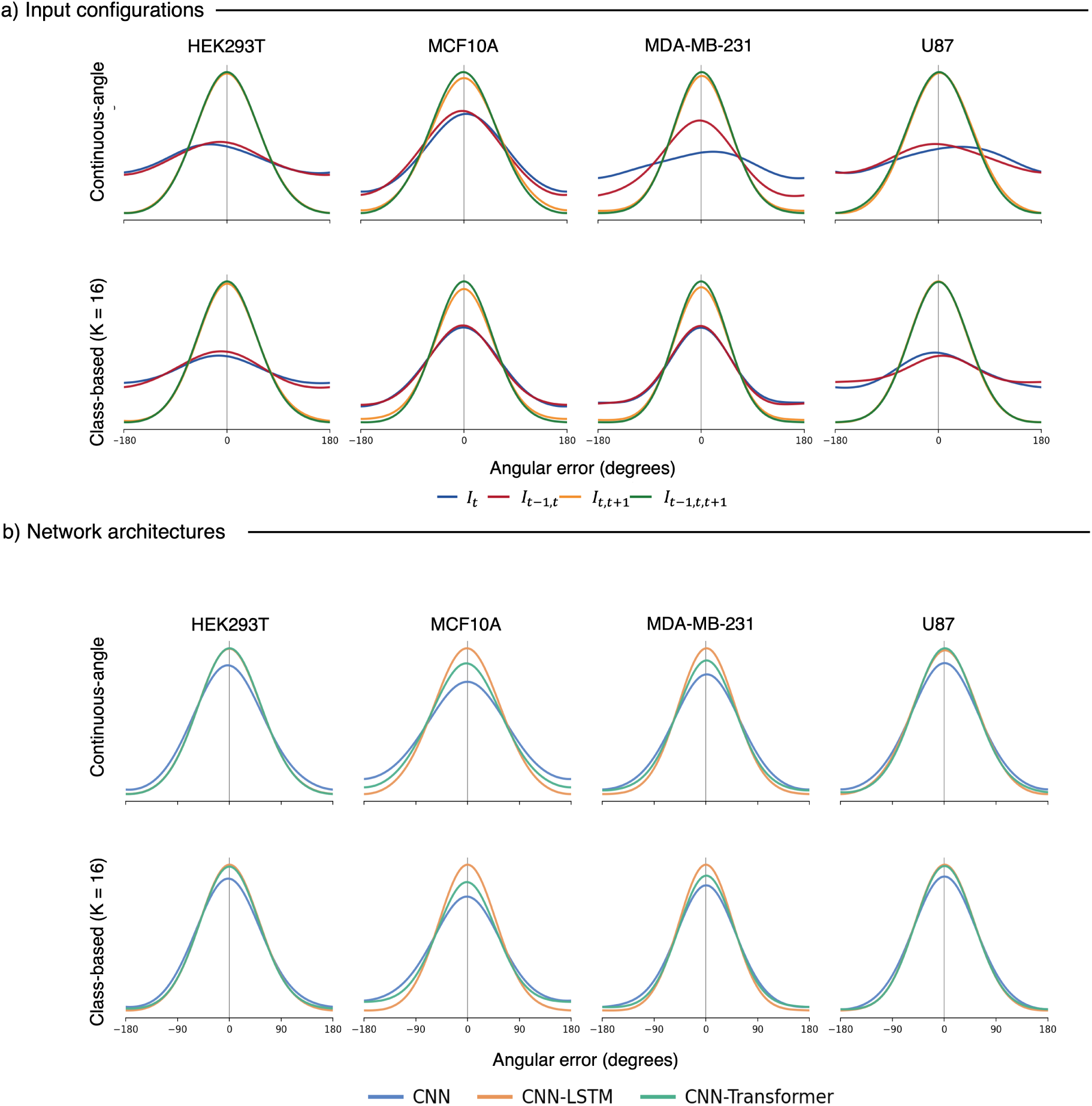
Angular error of the predicted direction at GAP 8. Each column is one cell line. The top row shows the continuous-angle models and the bottom row the class-based (*K* = 16) models. (a) **Input configurations.** Error distributions of the CNN–LSTM for the four image-patch configurations *I_t_*, *I_t−_*_1*,t*_, *I_t,t_*_+1_, and *I_t−_*_1*,t,t*+1_. (b) **Network architectures.** Error distributions of the CNN, CNN–LSTM, and CNN–Transformer with input *I_t−_*_1*,t,t*+1_. A narrower distribution centered at 0*^◦^* indicates more accurate direction predictions.

#### 4.3.3 The impact of model architecture

The previous section shows that direction prediction depends on which frames the patch contains. We next ask whether the way these frames are combined also matters. The CNN receives the frames together as input channels, while the CNN-LSTM and the CNN-Transformer receive them in temporal order. This comparison tests whether modelling the temporal order of the frames improves the prediction beyond providing the frames themselves. Figure 5b shows the angular-error distributions of the three architectures on the *I_t−_*_1*,t,t*+1_ input at GAP 8, for both representations. The CNN has the largest errors in every cell line. Averaged over the four cell lines, its median angular error is 36.4*^◦^* for the continuous-angle representation and 29.3*^◦^* for the class-based representation, compared with 25.3*^◦^* and 18.1*^◦^* for the CNN-LSTM and 27.3*^◦^* and 20.8*^◦^* for the CNN-Transformer. The two temporal models differ by less than 3*^◦^* on average, and the CNN-LSTM has the lowest error in every cell line except U87 with the continuous-angle representation.

#### 4.3.4 Velocity prediction

The previous sections show how the predicted direction depends on the input patches and the model architecture. The predicted position of a cell also depends on its displacement, which is not known at test time. We next ask how the choice of displacement estimation affects linking performance. Figure 6 compares each method described in Section 3.4 with the distance-only LAP baseline, for both representations.

**Figure 6:**
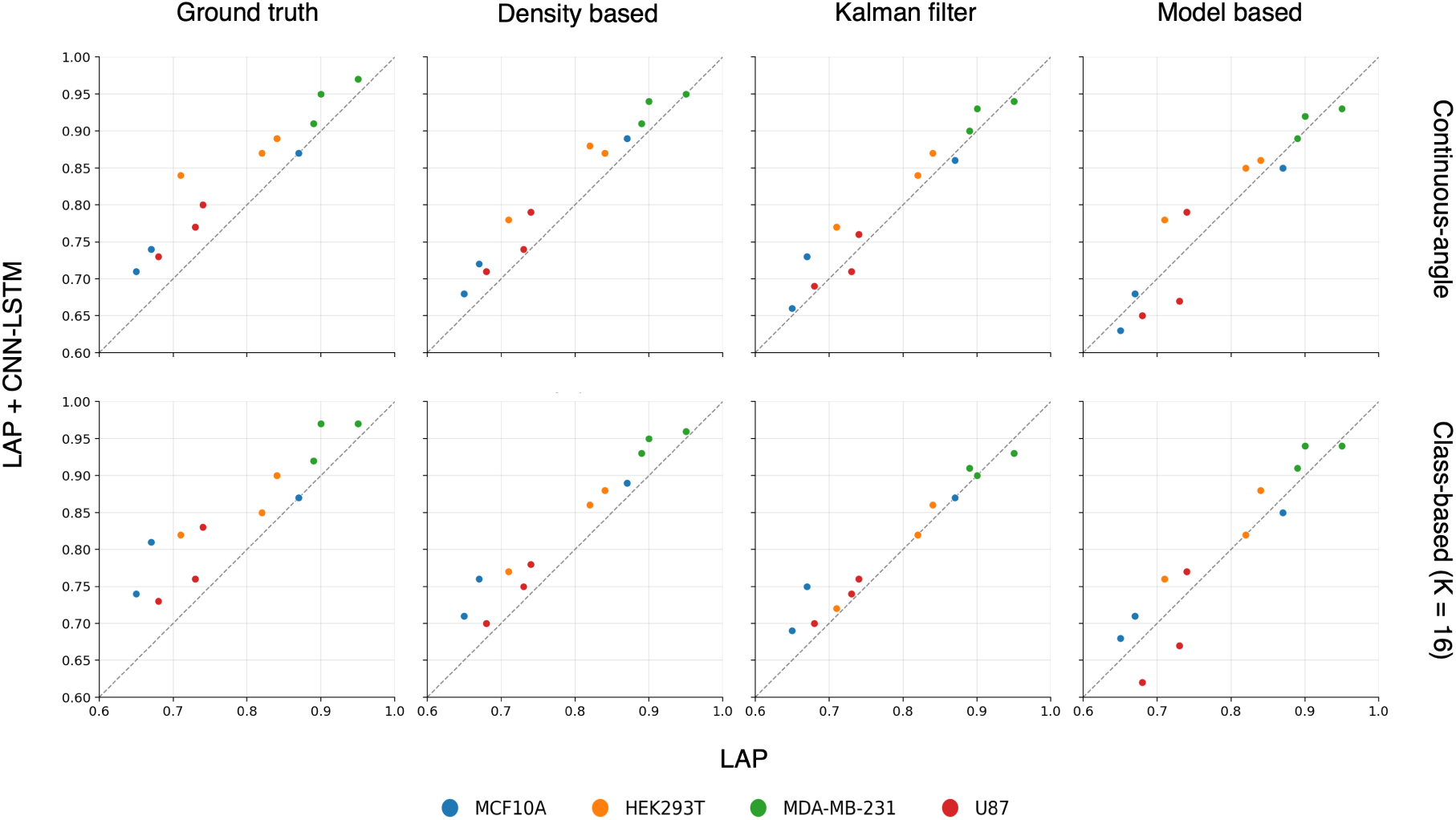
Linking performance at GAP 8 with the direction predicted by the CNN-LSTM, one column per displacement estimation method. The top row shows the continuous-angle models and the bottom row the class-based (*K* = 16) models. Each dot is one sequence, coloured by cell line. The x-axis gives the linking performance of the distance-only LAP baseline and the y-axis the linking performance of LAP with the predicted direction. The dashed line marks equal performance, so dots above the line show an improvement over the baseline and dots below it a decrease.

With the ground-truth displacement, the predicted direction increases linking performance by 5.0 percentage points on average for the continuous-angle representation and by 6.0 points for the classbased representation. With the model-based displacement, the improvement drops to 0.4 and 0.8 points, and linking performance is lower than the baseline in 5 and 4 of the 12 sequences. Without the ground-truth displacement, the density based displacement gives the largest improvement: 3.4 points for the continuous-angle representation and 4.1 points for the class-based representation, with no sequence below the baseline. The Kalman filter improves linking performance by 1.8 and 1.7 points.

## 5 Conclusion

In this work we studied whether deep-learning models can predict the movement direction of a cell from image patches, and whether this prediction improves cell linking in low-frame-rate microscopy. Our results show that the predicted direction improves linking compared with the distance-only LAP baseline at GAP 8, and that most of the useful information comes from the target frame. With a single image *I_t_*, the errors are spread over almost all directions for most cell lines, so the appearance of the cell at one time point is not enough to predict its movement over two hours. Adding past information with *I_t−_*_1*,t*_ improves the prediction only slightly for most cell lines, while adding the future patch with *I_t,t_*_+1_ or *I_t−_*_1*,t,t*+1_ concentrates the predictions close to the true direction. The future patch contains no marker identifying the cell, but it provides visual information that helps the model infer its movement direction between frames.

The accuracy of the predictions also depends on how the direction is represented. Previous work has predicted the movement direction either as one of four direction classes [39] or as a continuous angle [9], and the continuous angle was more accurate than the four classes. In our experiments, with *K* = 16 classes, the class-based models have lower angular errors than the continuous-angle models for all cell lines.

Beyond the direction representation, prediction accuracy also depends on the model architecture. With the same input, the CNN-LSTM gave the lowest angular errors on average, followed closely by the CNN-Transformer, while the CNN had the largest errors in every cell line. The best-performing configuration therefore combines the CNN-LSTM with the class-based representation.

Even with the best models, the predicted directions are not exact: the median angular error is between 9*^◦^* and 33*^◦^*. The controlled analysis in Section 4.2 shows that linking performance changes little for noise levels up to *σ* = 20–30*^◦^*, which correspond to median angular errors of about 14–21*^◦^*. With the density based displacement, these predictions still improve linking compared with the distance-only LAP baseline, by 3.4 percentage points with the continuous-angle representation and by 4.1 points with the class-based representation. In the controlled analysis, both representations give similar linking performance at the same noise level. This suggests that the larger improvement with the class-based representation comes from its more accurate direction predictions.

## Acknowledgments

The authors gratefully acknowledge the support provided by the National Institute Of General Medical Sciences of the National Institutes of Health under Award Number R21GM150066 and R35GM141891. Results were generated using the Palmetto Cluster at Clemson University, supported by the National Science Foundation under Grant numbers MRI 1228312, II NEW 1405767, MRI 1725573, and MRI 2018069. Any opinions, findings, and conclusions or recommendations expressed in this material are those of the author(s) and do not necessarily reflect the views of the funding agencies.

## Author Contributions

M.G. developed the code, trained the models, performed the experiments and analysis, and led manuscript writing. B.N. contributed to the experimental comparison with other tracking methods. T.B. acquired the imaging data. M.R.B. and F.I. secured funding and contributed to manuscript writing. All authors reviewed and approved the final manuscript.

## Supplemental Information

The main text reports the results of CNN-LSTM at GAP 8. Here, we report the same analyses for CNN, CNN-LSTM, and CNN-Transformer at GAP 1, 2, 4, and 8. Sections S1, S2, and S3 follow the order of the main text: angular-error distributions, predicting direction from image patches, and velocity prediction.

### S1 Angular-error distributions

**Figure S1:**
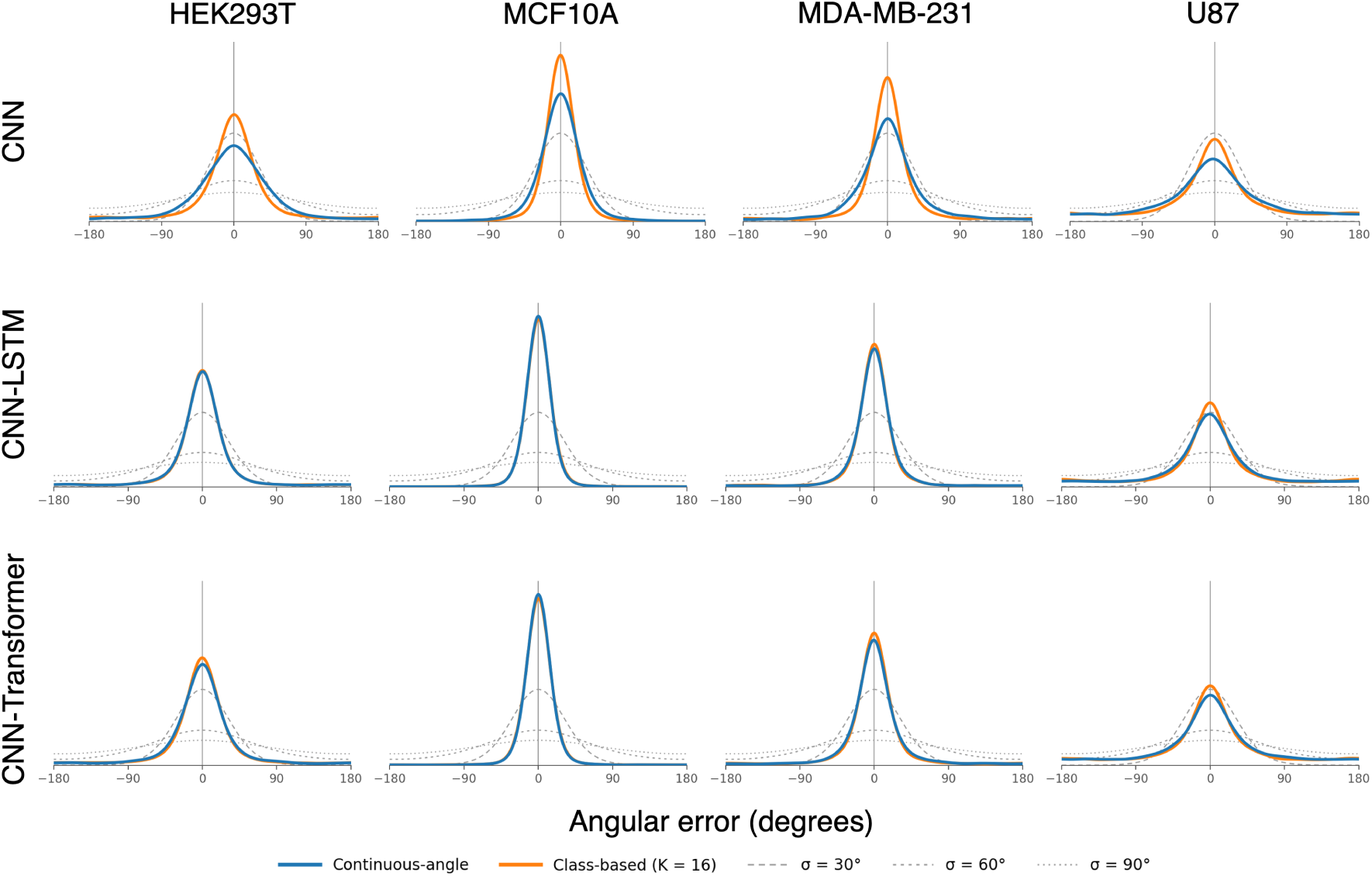
Angular error distributions at GAP 1 for the continuous-angle and class-based (*K* = 16) representations with input *I_t−_*_1*,t,t*+1_, related to Figure 4. Each row is one architecture (CNN, CNN–LSTM, and CNN–Transformer) and each column is one cell line. Von Mises reference distributions at *σ* = 30*^◦^*, 60*^◦^*, and 90*^◦^* are shown in grey.

**Figure S2:**
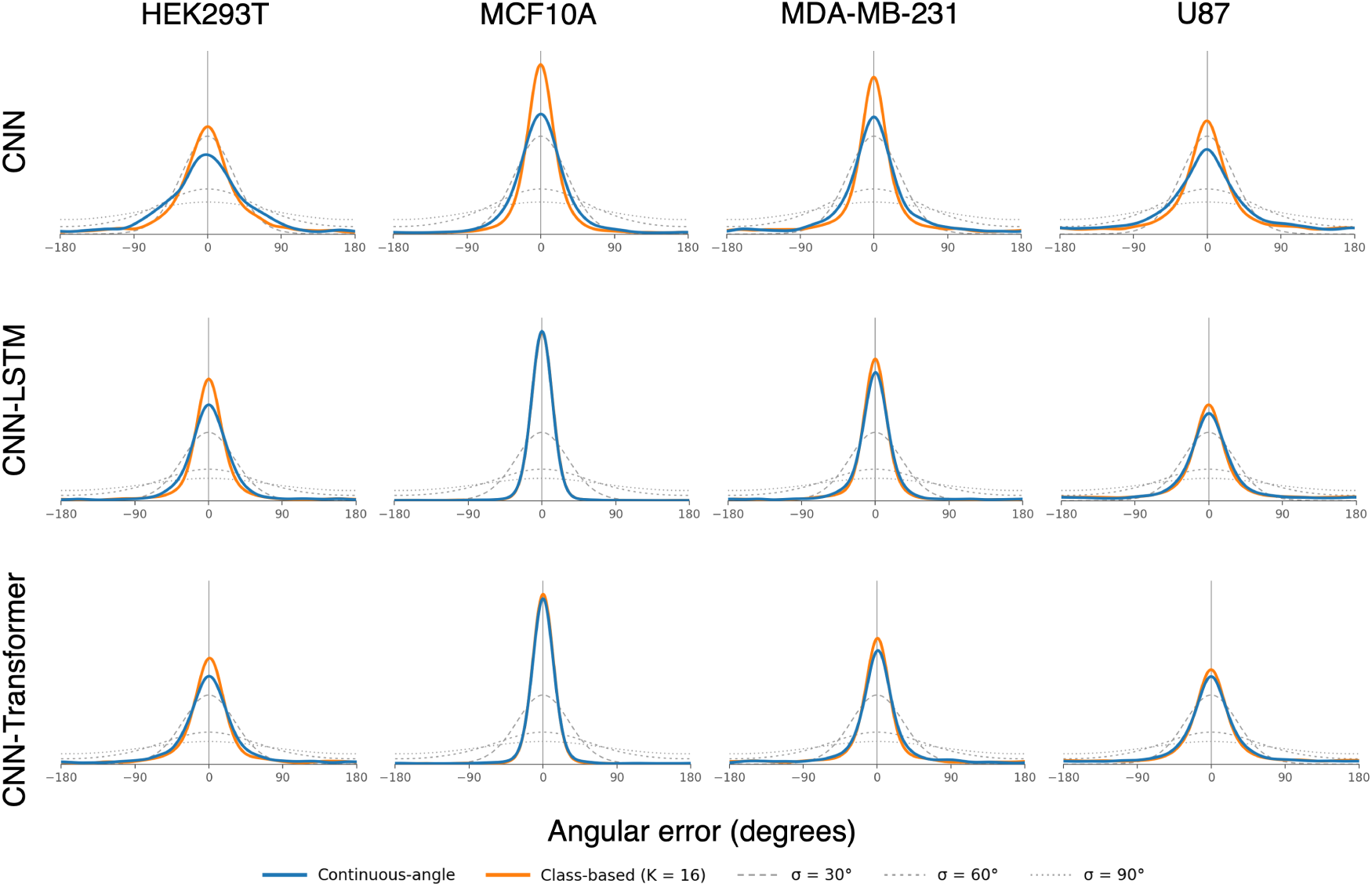
Angular error distributions at GAP 2. Panels as in Figure S1.

**Figure S3:**
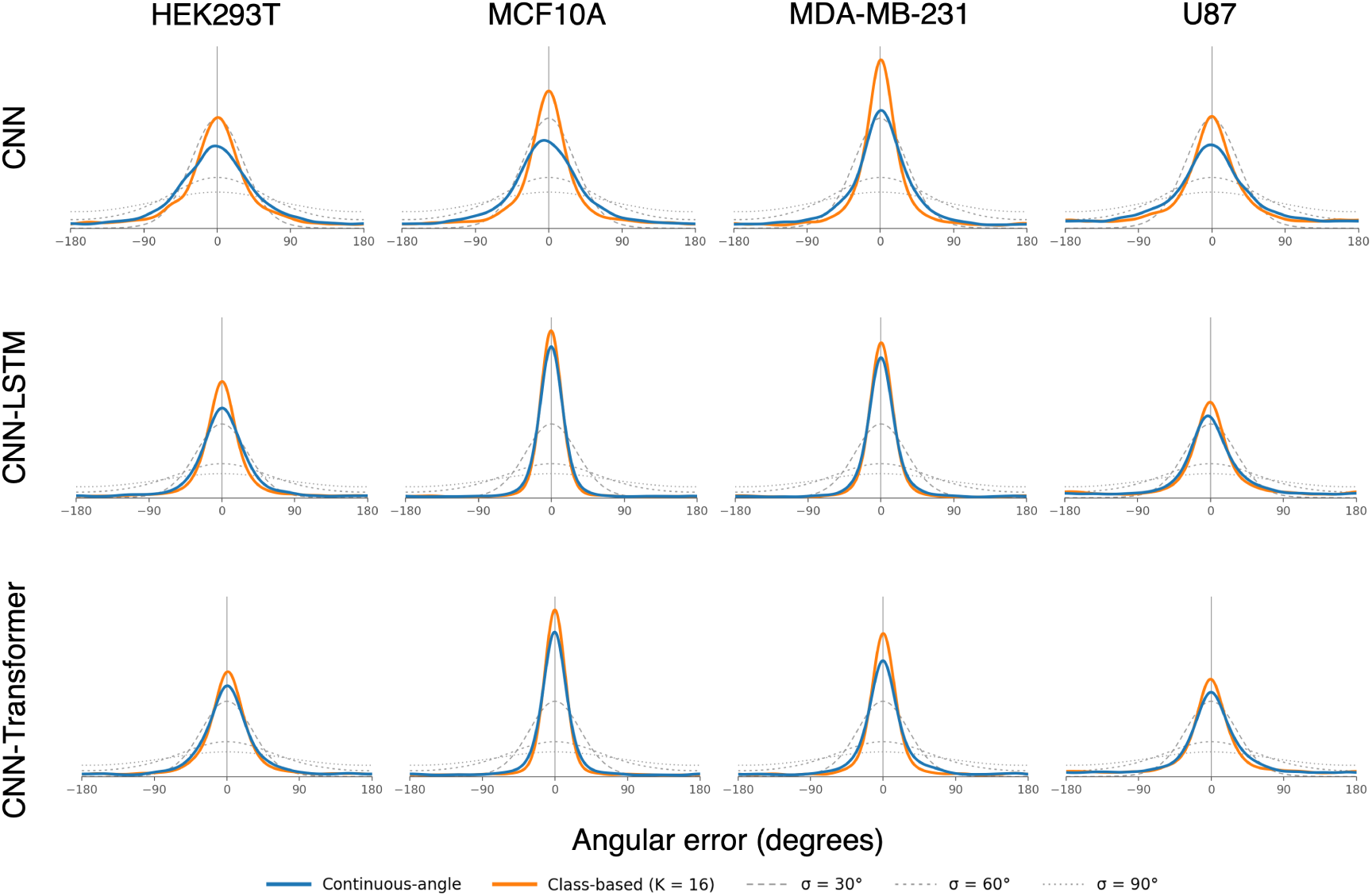
Angular error distributions at GAP 4. Panels as in Figure S1.

**Figure S4:**
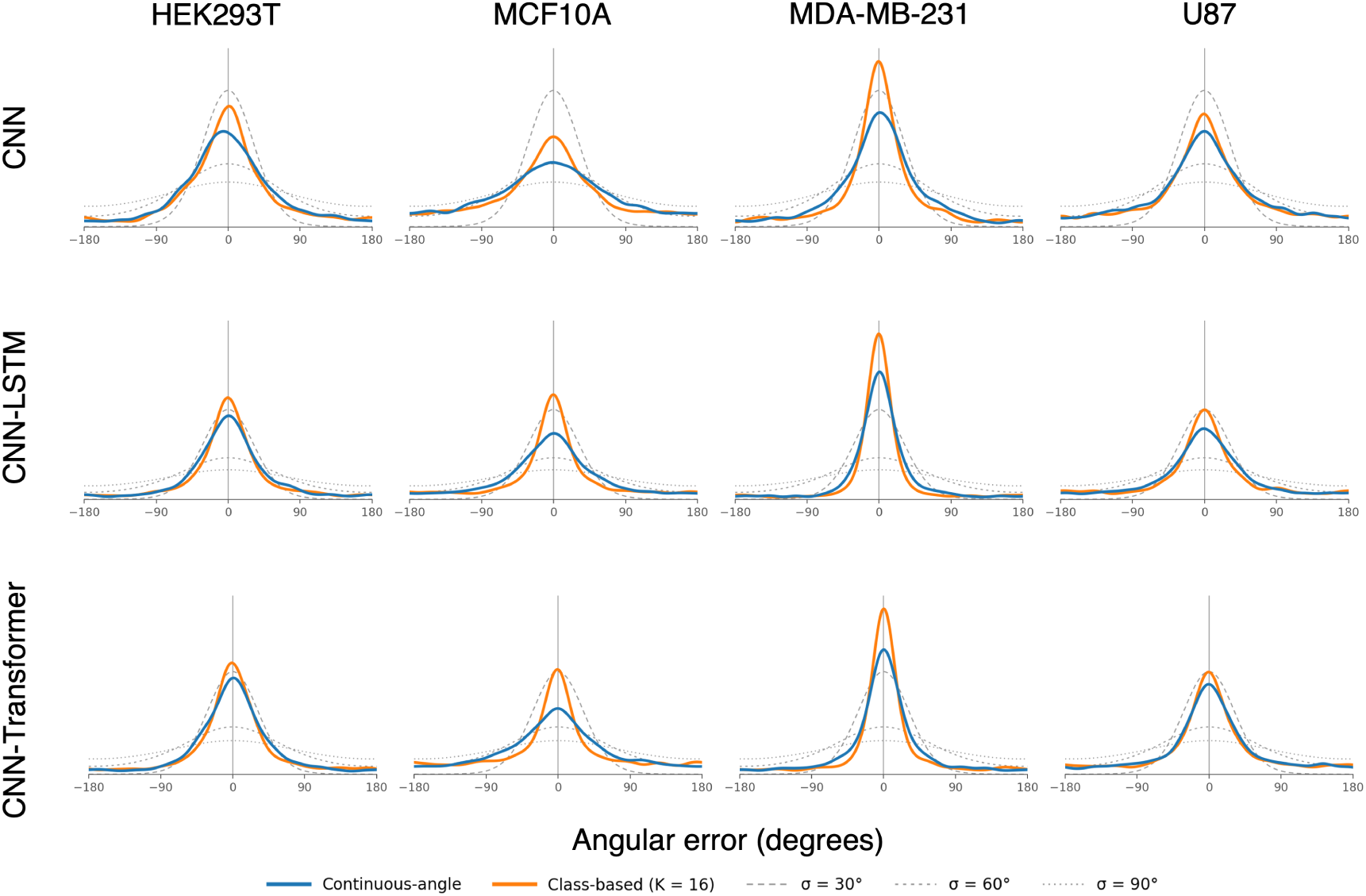
Angular error distributions at GAP 8. Panels as in Figure S1.

### S2 Predicting direction from image patches

#### S2.1 Class-based

**Figure S5:**
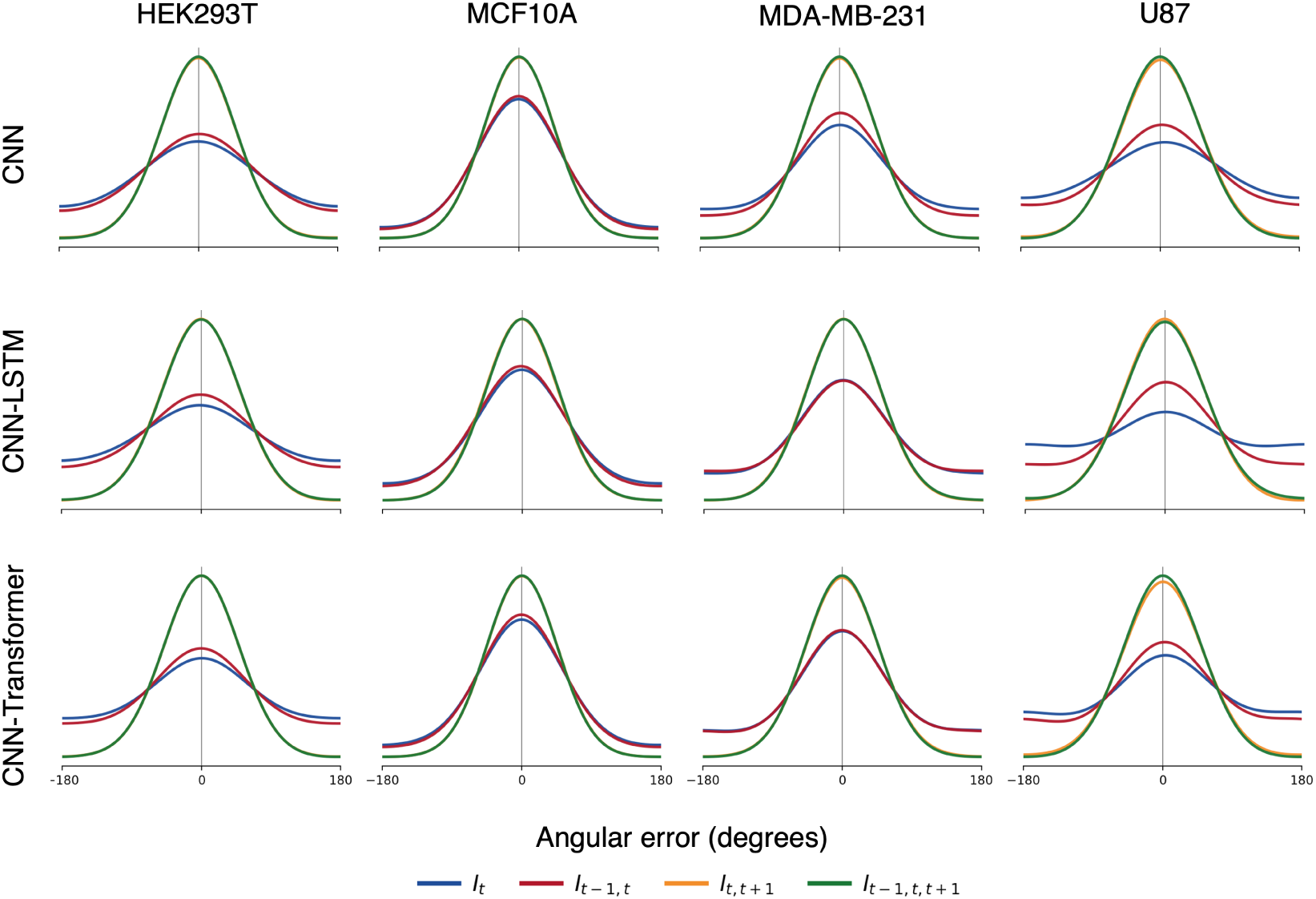
Angular error of the class-based (*K* = 16) direction predictions at GAP 1 for the four image-patch configurations, related to Figure 5. Each row is one architecture (CNN, CNN–LSTM, and CNN–Transformer) and each column is one cell line.

**Figure S6:**
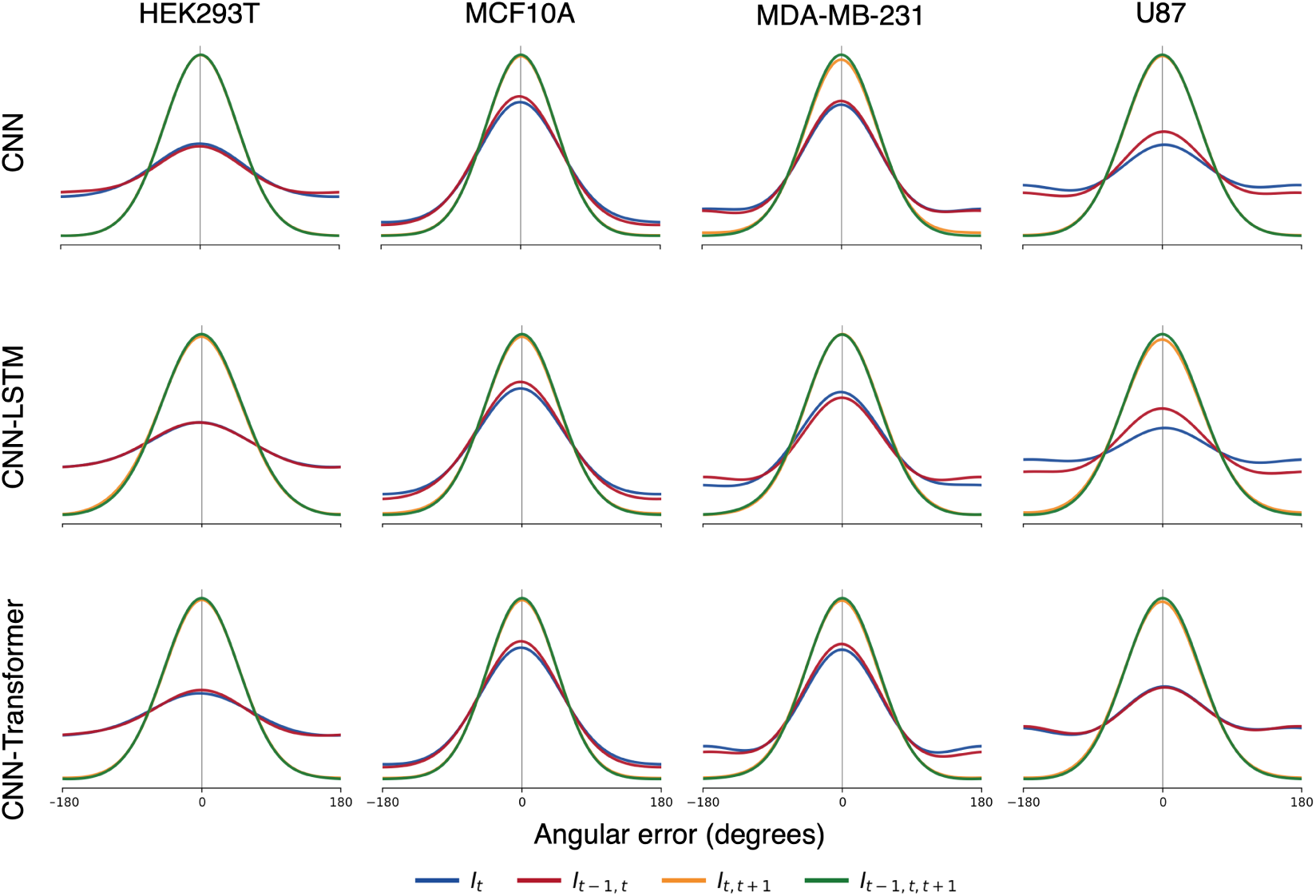
Angular error of the class-based direction predictions at GAP 2. Panels as in Figure S5.

**Figure S7:**
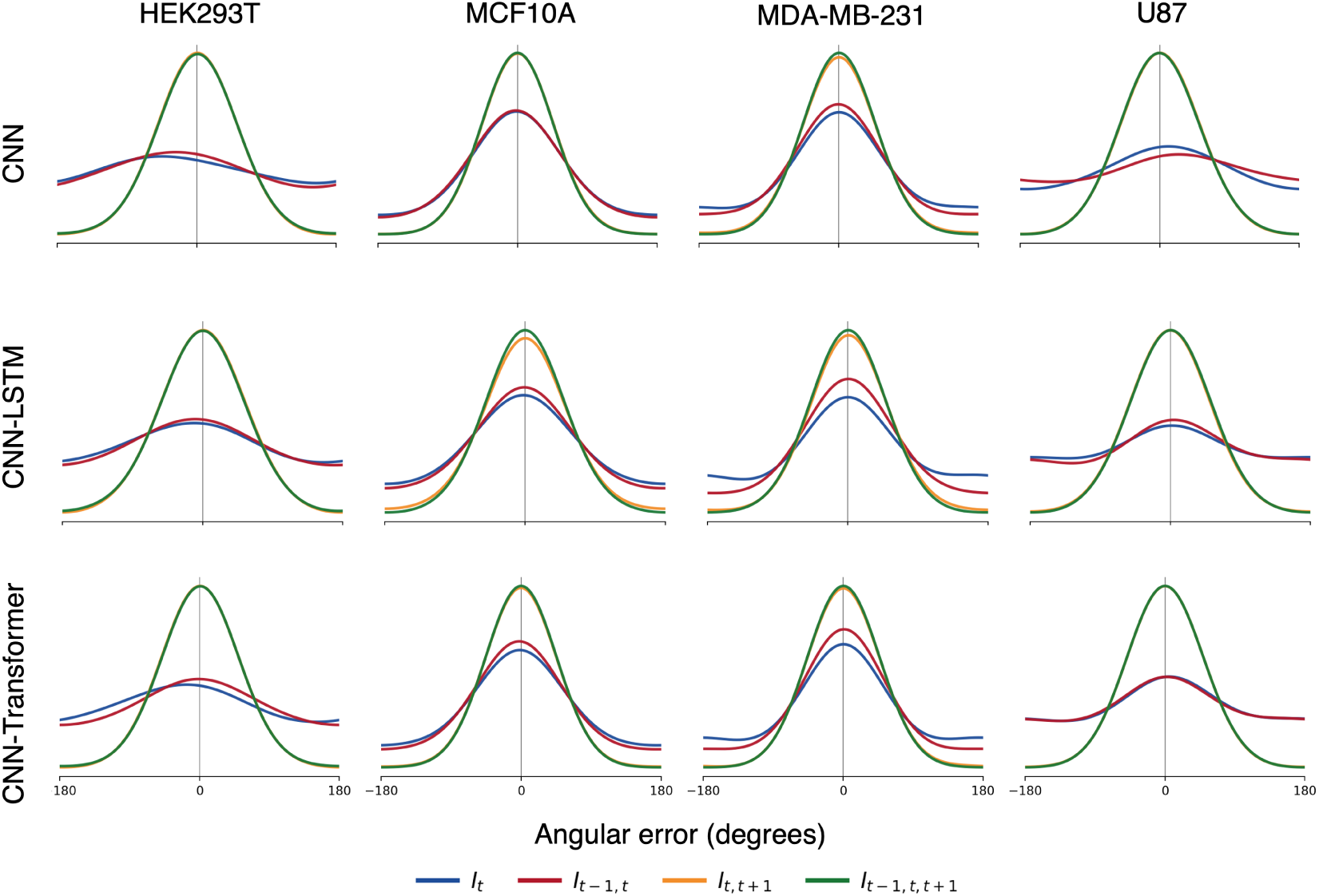
Angular error of the class-based direction predictions at GAP 4. Panels as in Figure S5.

**Figure S8:**
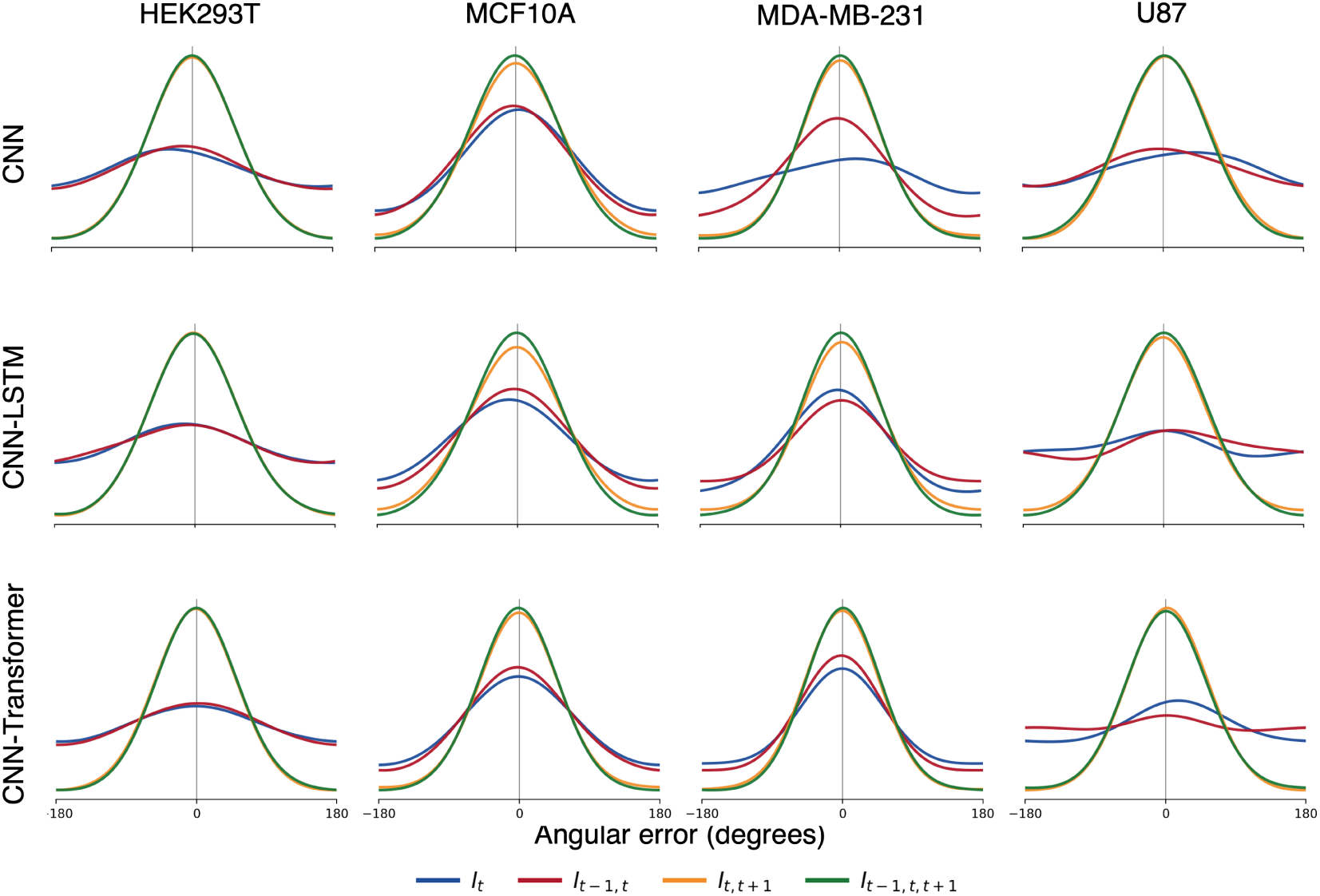
Angular error of the class-based direction predictions at GAP 8. Panels as in Figure S5.

#### S2.2 Continuous-angle

**Figure S9:**
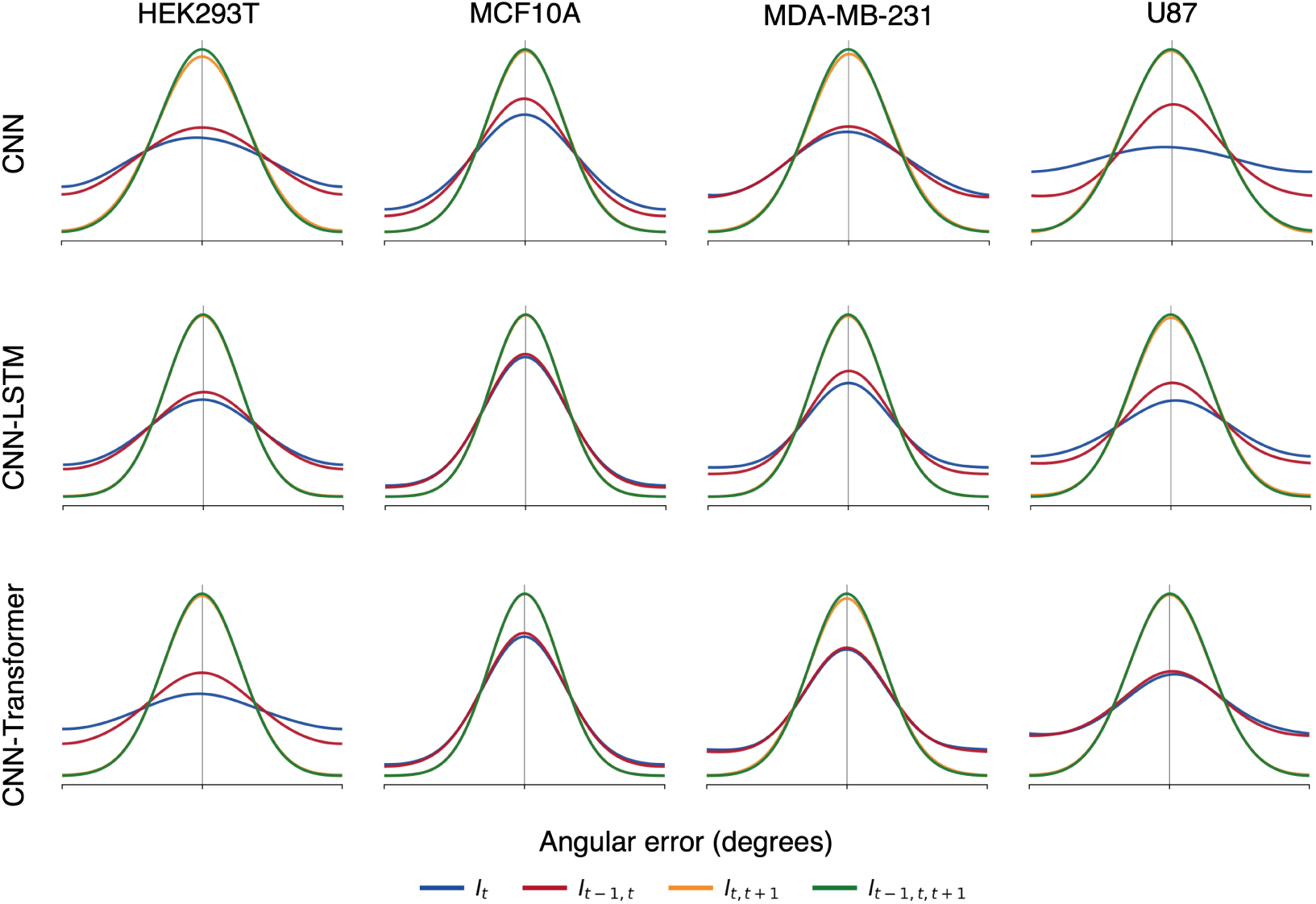
Angular error of the continuous-angle direction predictions at GAP 1 for the four image-patch configurations, related to Figure 5. Each row is one architecture (CNN, CNN–LSTM, and CNN–Transformer) and each column is one cell line.

**Figure S10:**
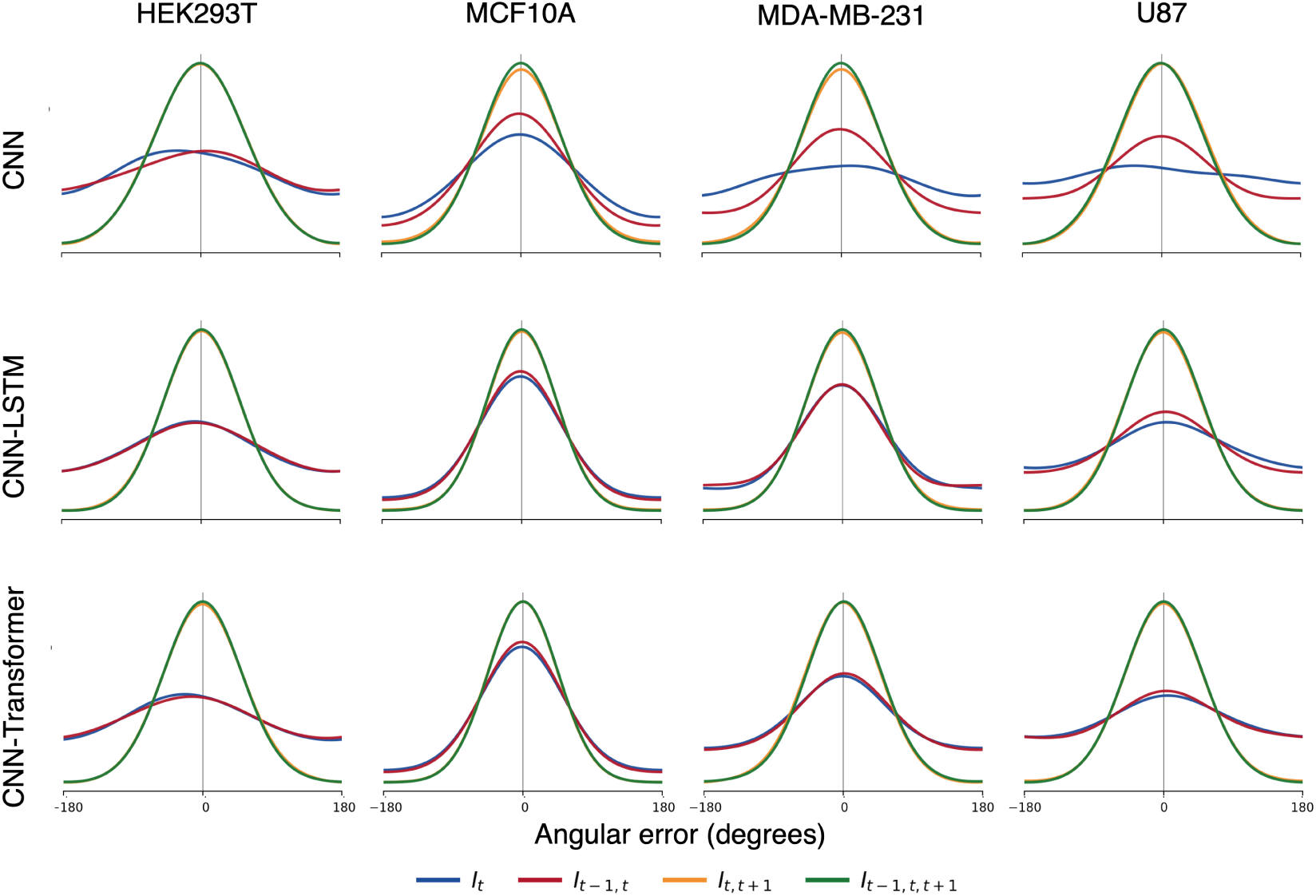
Angular error of the continuous-angle direction predictions at GAP 2. Panels as in Figure S9.

**Figure S11:**
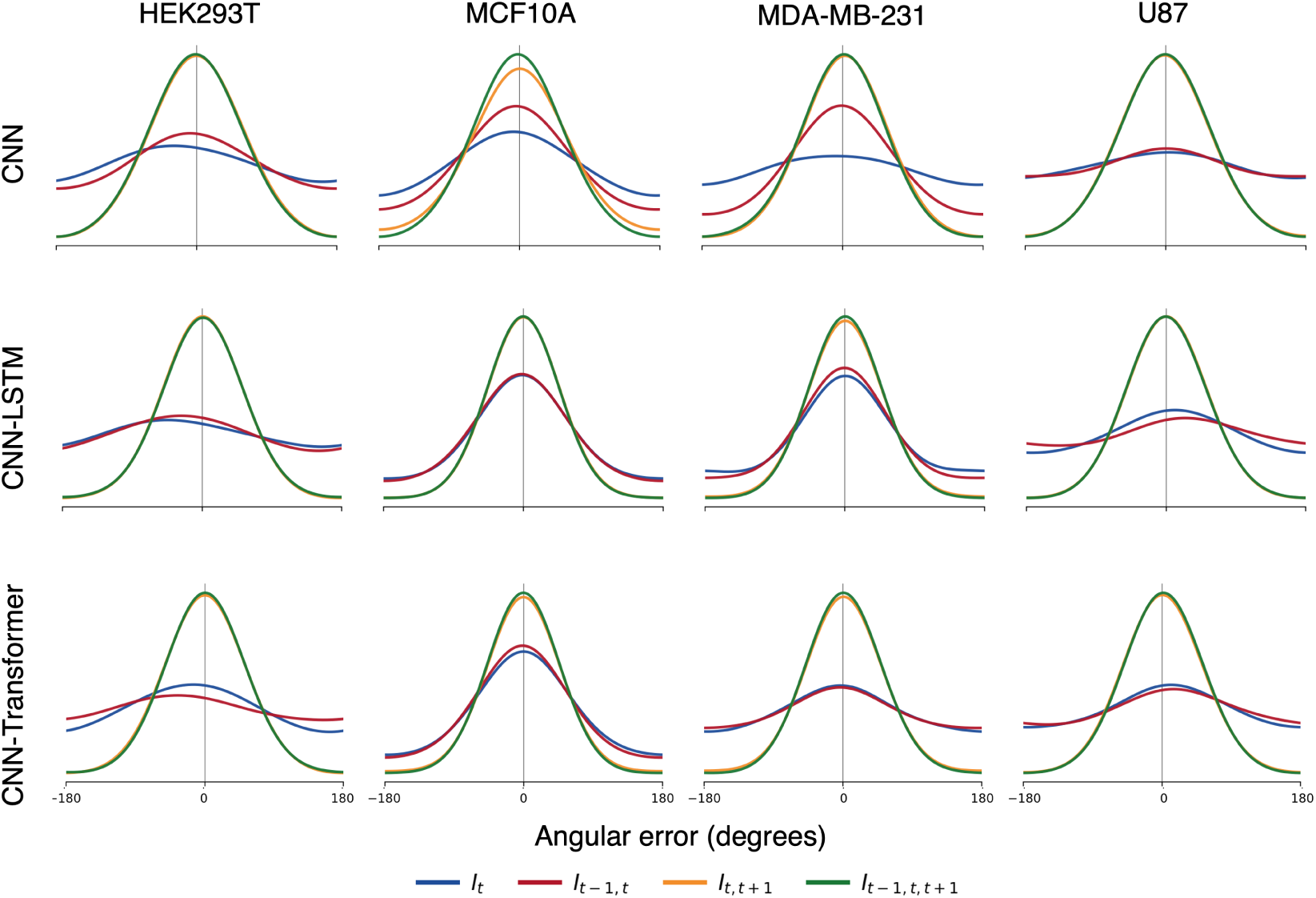
Angular error of the continuous-angle direction predictions at GAP 4. Panels as in Figure S9.

**Figure S12:**
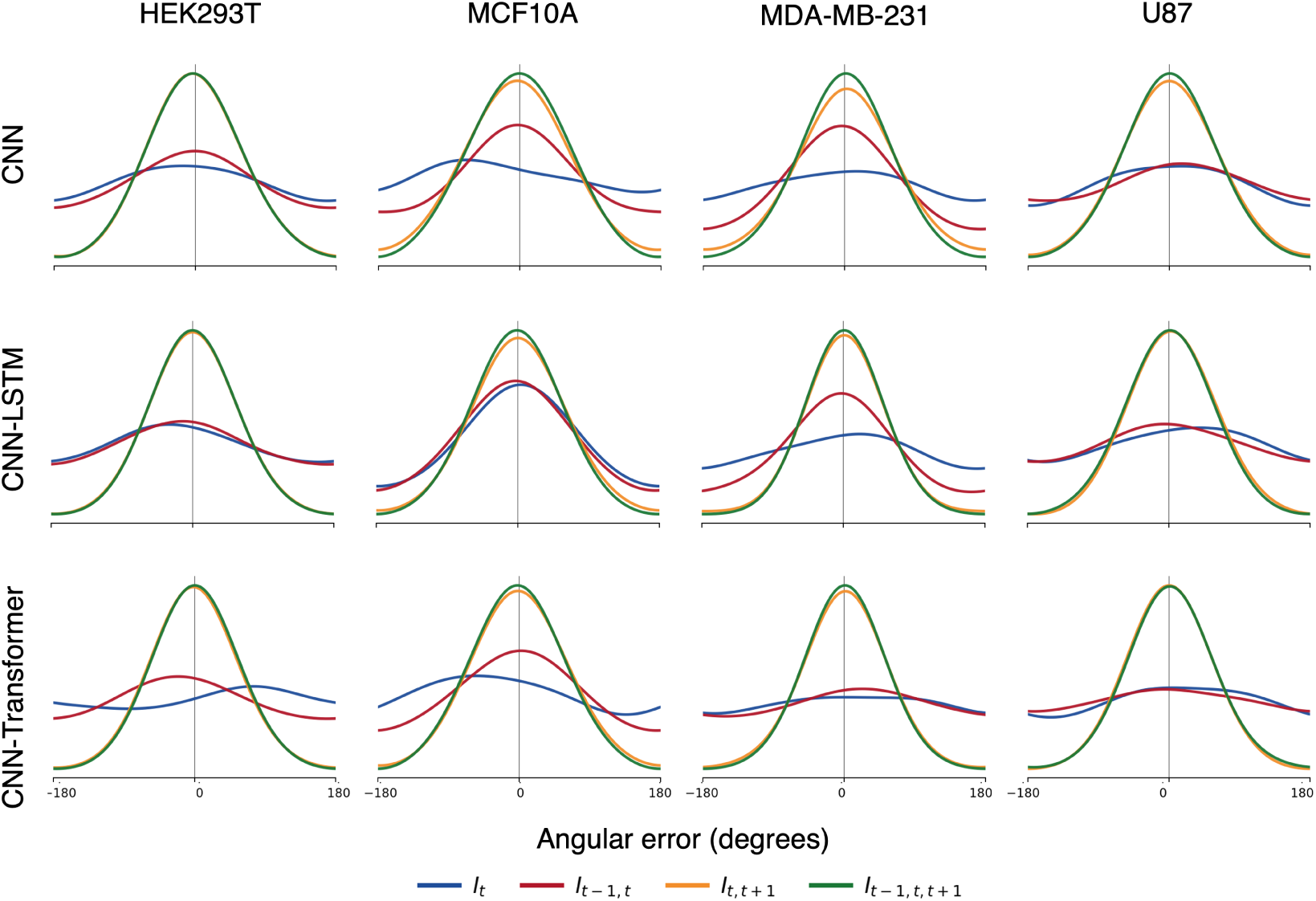
Angular error of the continuous-angle direction predictions at GAP 8. Panels as in Figure S9.

#### S2.3 Network architectures

**Figure S13:**
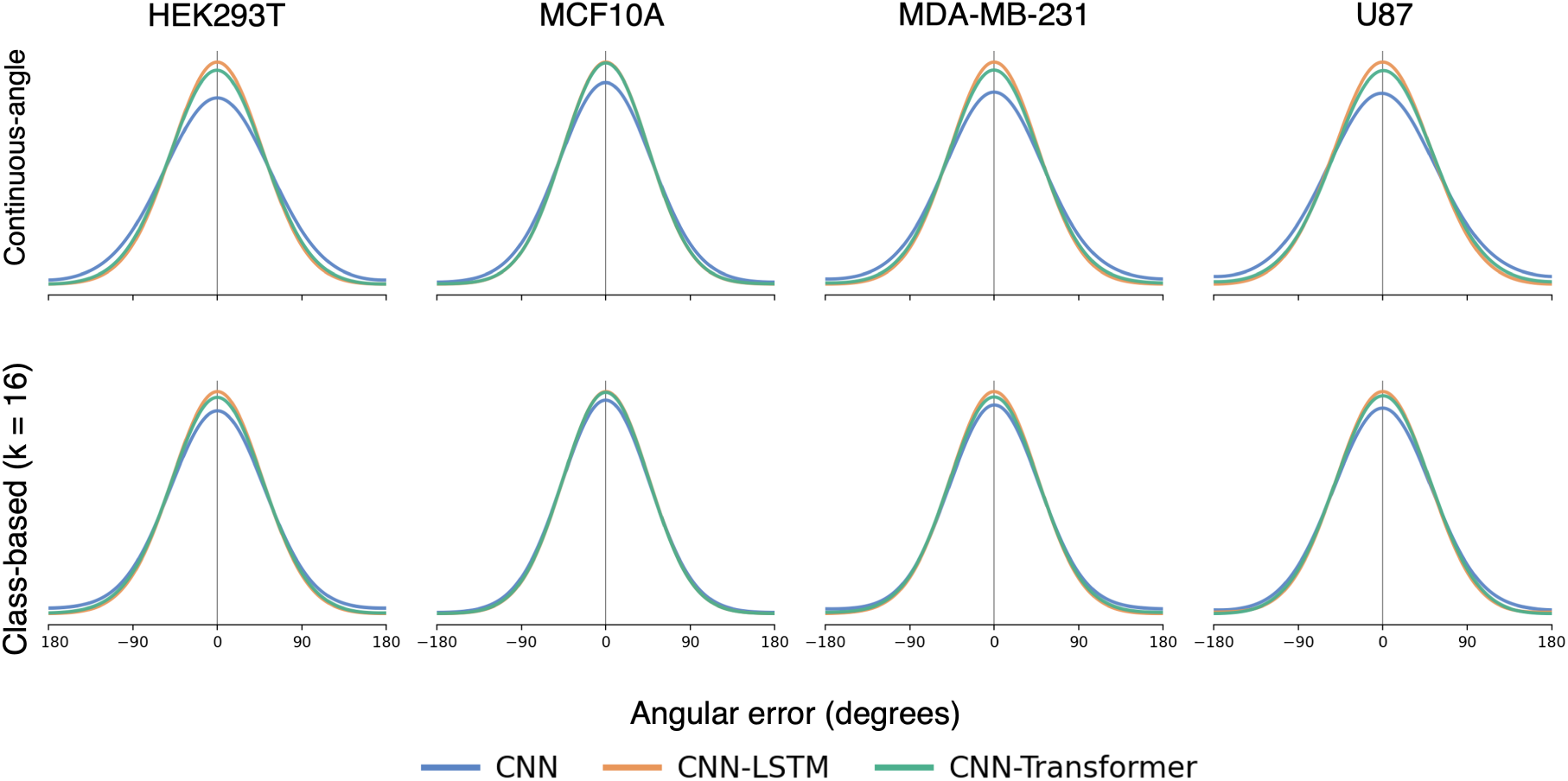
Network architectures at GAP 1, related to Figure 5. Error distributions of the CNN, CNN–LSTM, and CNN–Transformer with input *I_t−_*_1*,t,t*+1_. A narrower distribution centered at 0*^◦^* indicates more accurate direction predictions.

**Figure S14:**
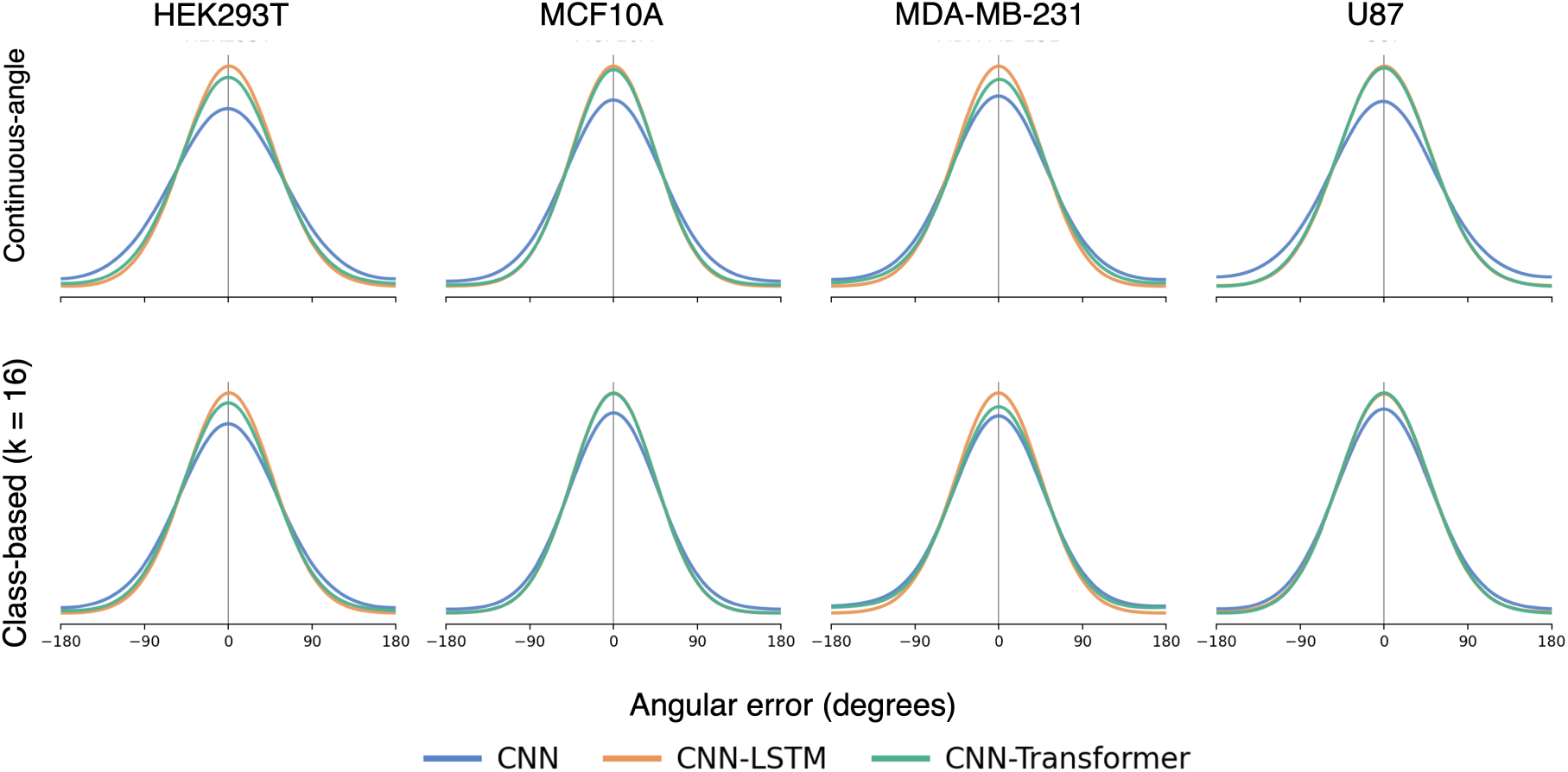
Network architectures at GAP 2. Error distributions of the CNN, CNN–LSTM, and CNN–Transformer with input *I_t−_*_1*,t,t*+1_. A narrower distribution centered at 0*^◦^* indicates more accurate direction predictions.

**Figure S15:**
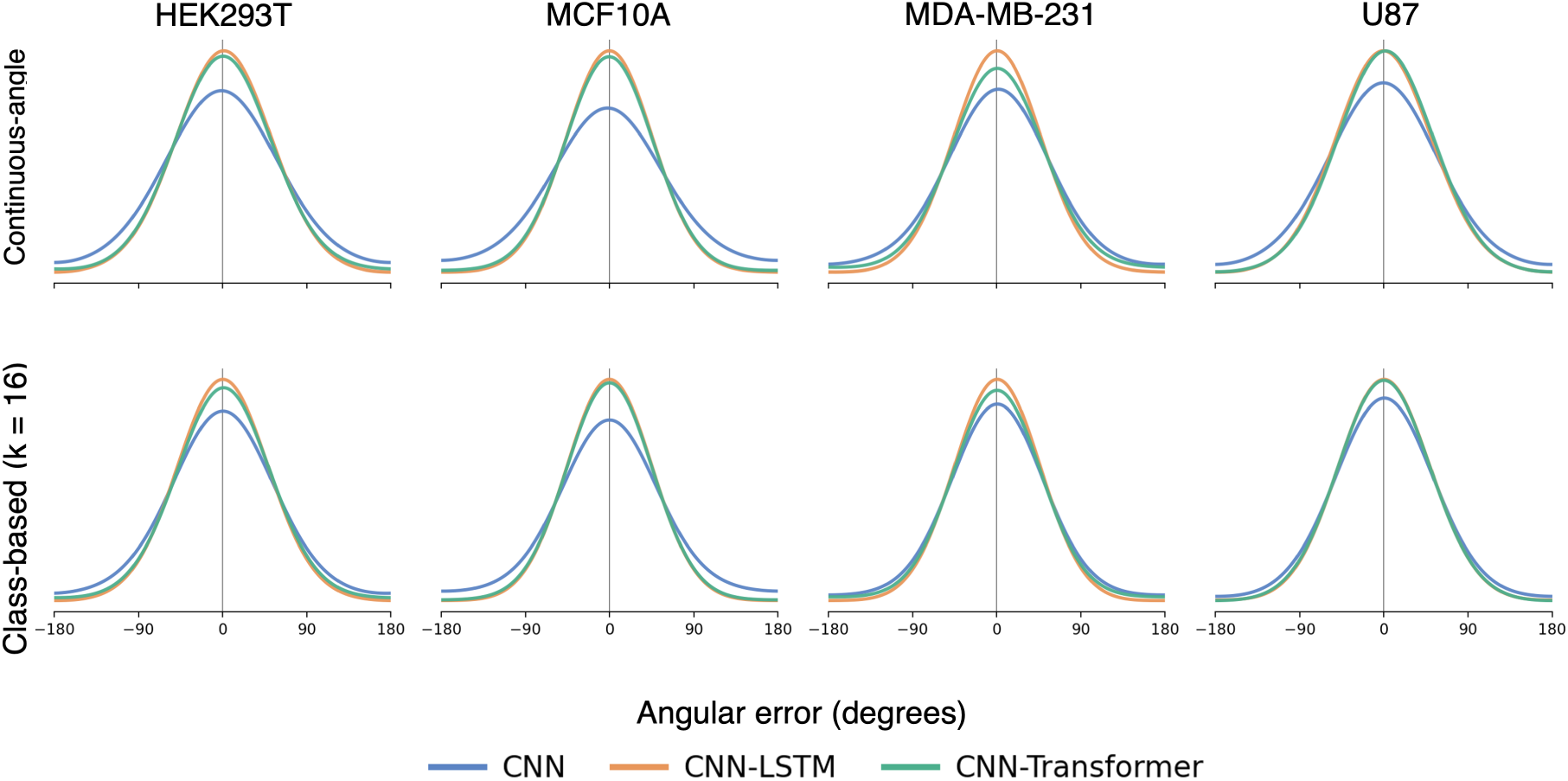
Network architectures at GAP 4. Error distributions of the CNN, CNN–LSTM, and CNN–Transformer with input *I_t−_*_1*,t,t*+1_. A narrower distribution centered at 0*^◦^* indicates more accurate direction predictions.

### S3 Velocity prediction

**Table S1:** Linking performance at GAP 1 and GAP 2, related to Figure 6. Values are the average recall over the test sequences of each cell line, for the distance-only LAP baseline and for LAP with the direction predicted by each architecture, for the four displacement estimation methods.

|  |  | Ground truth |  |  | Density based |  |  | Kalman filter |  |  | Model based |  |  |
|---|---|---|---|---|---|---|---|---|---|---|---|---|---|
| Cell line | LAP | CNN | LSTM | Trans. | CNN | LSTM | Trans. | CNN | LSTM | Trans. | CNN | LSTM | Trans. |
| **GAP 1** |  |  |  |  |  |  |  |  |  |  |  |  |  |
| *Class-based* (*K* = 16) |  |  |  |  |  |  |  |  |  |  |  |  |  |
| HEK293T | 0.99 | 0.99 | 0.99 | 0.99 | 0.99 | 0.99 | 0.99 | 0.99 | 0.99 | 0.99 | 0.99 | 0.99 | 0.99 |
| MCF10A | 0.99 | 0.99 | 0.99 | 0.99 | 0.99 | 0.99 | 0.99 | 0.99 | 0.99 | 0.99 | 0.99 | 0.99 | 0.99 |
| MDA-MB-231 | 0.99 | 0.99 | 0.99 | 0.99 | 0.99 | 0.99 | 0.99 | 0.99 | 0.99 | 0.99 | 0.99 | 0.99 | 0.99 |
| U87 | 0.98 | 0.97 | 0.98 | 0.97 | 0.98 | 0.98 | 0.98 | 0.97 | 0.97 | 0.97 | 0.97 | 0.97 | 0.97 |
| *Continuous-angle* |  |  |  |  |  |  |  |  |  |  |  |  |  |
| HEK293T | 0.99 | 0.99 | 0.99 | 0.99 | 0.99 | 0.99 | 0.99 | 0.99 | 0.99 | 0.99 | 0.99 | 0.99 | 0.99 |
| MCF10A | 0.99 | 0.99 | 0.99 | 0.99 | 0.99 | 0.99 | 0.99 | 0.99 | 0.99 | 0.99 | 0.99 | 0.99 | 0.99 |
| MDA-MB-231 | 0.99 | 0.99 | 0.99 | 0.99 | 0.99 | 0.99 | 0.99 | 0.99 | 0.99 | 0.99 | 0.99 | 0.99 | 0.99 |
| U87 | 0.98 | 0.96 | 0.97 | 0.96 | 0.98 | 0.98 | 0.98 | 0.98 | 0.98 | 0.98 | 0.97 | 0.97 | 0.97 |
| **GAP 2** |  |  |  |  |  |  |  |  |  |  |  |  |  |
| *Class-based* (*K* = 16) |  |  |  |  |  |  |  |  |  |  |  |  |  |
| HEK293T | 0.98 | 0.97 | 0.98 | 0.98 | 0.99 | 0.99 | 0.99 | 0.97 | 0.98 | 0.98 | 0.97 | 0.98 | 0.98 |
| MCF10A | 0.99 | 0.98 | 0.99 | 0.99 | 0.99 | 0.99 | 0.99 | 0.99 | 0.99 | 0.99 | 0.98 | 0.99 | 0.99 |
| MDA-MB-231 | 0.98 | 0.98 | 0.99 | 0.99 | 0.99 | 0.99 | 0.99 | 0.99 | 0.99 | 0.99 | 0.98 | 0.98 | 0.98 |
| U87 | 0.92 | 0.92 | 0.93 | 0.93 | 0.93 | 0.93 | 0.93 | 0.92 | 0.92 | 0.92 | 0.89 | 0.91 | 0.91 |
| *Continuous-angle* |  |  |  |  |  |  |  |  |  |  |  |  |  |
| HEK293T | 0.98 | 0.97 | 0.98 | 0.98 | 0.98 | 0.98 | 0.98 | 0.97 | 0.98 | 0.98 | 0.97 | 0.98 | 0.98 |
| MCF10A | 0.99 | 0.98 | 0.99 | 0.99 | 0.99 | 0.99 | 0.99 | 0.99 | 0.99 | 0.99 | 0.98 | 0.99 | 0.98 |
| MDA-MB-231 | 0.98 | 0.99 | 0.99 | 0.99 | 0.99 | 0.99 | 0.99 | 0.99 | 0.99 | 0.99 | 0.99 | 0.99 | 0.99 |
| U87 | 0.92 | 0.91 | 0.93 | 0.92 | 0.93 | 0.93 | 0.93 | 0.91 | 0.93 | 0.92 | 0.90 | 0.91 | 0.91 |

LSTM: CNN–LSTM; Trans.: CNN–Transformer.

#### S3.1 Class-based

**Figure S16:**
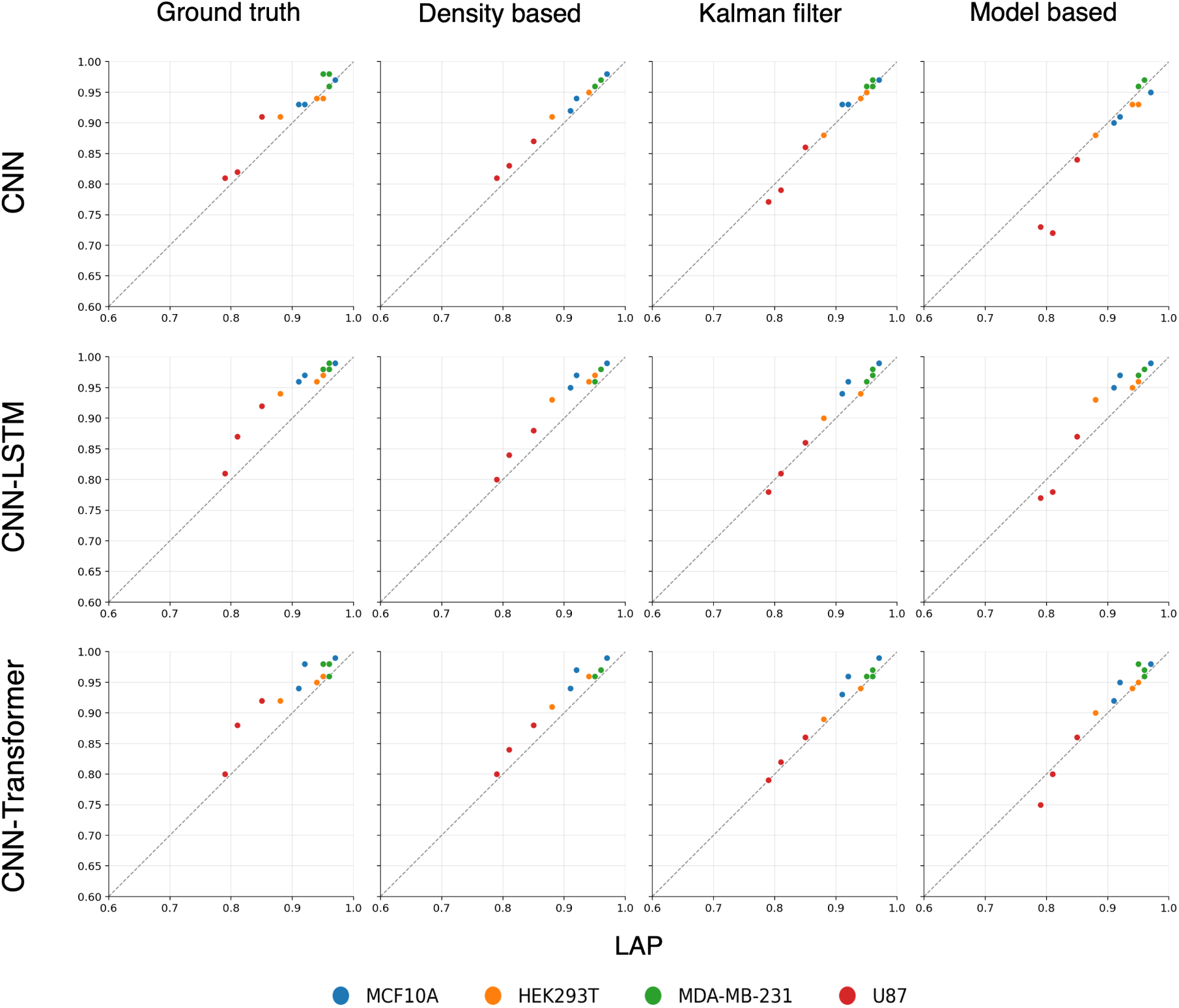
Linking performance of the class-based (*K* = 16) models at GAP 4, related to Figure 6. Each row is one architecture (CNN, CNN–LSTM, and CNN–Transformer) and each column is one displacement estimation method. Each dot is one test sequence, coloured by cell line. The x-axis gives the recall of the distance-only LAP baseline and the y-axis the recall of LAP with the predicted direction. Dots above the dashed line show an improvement over the baseline.

**Figure S17:**
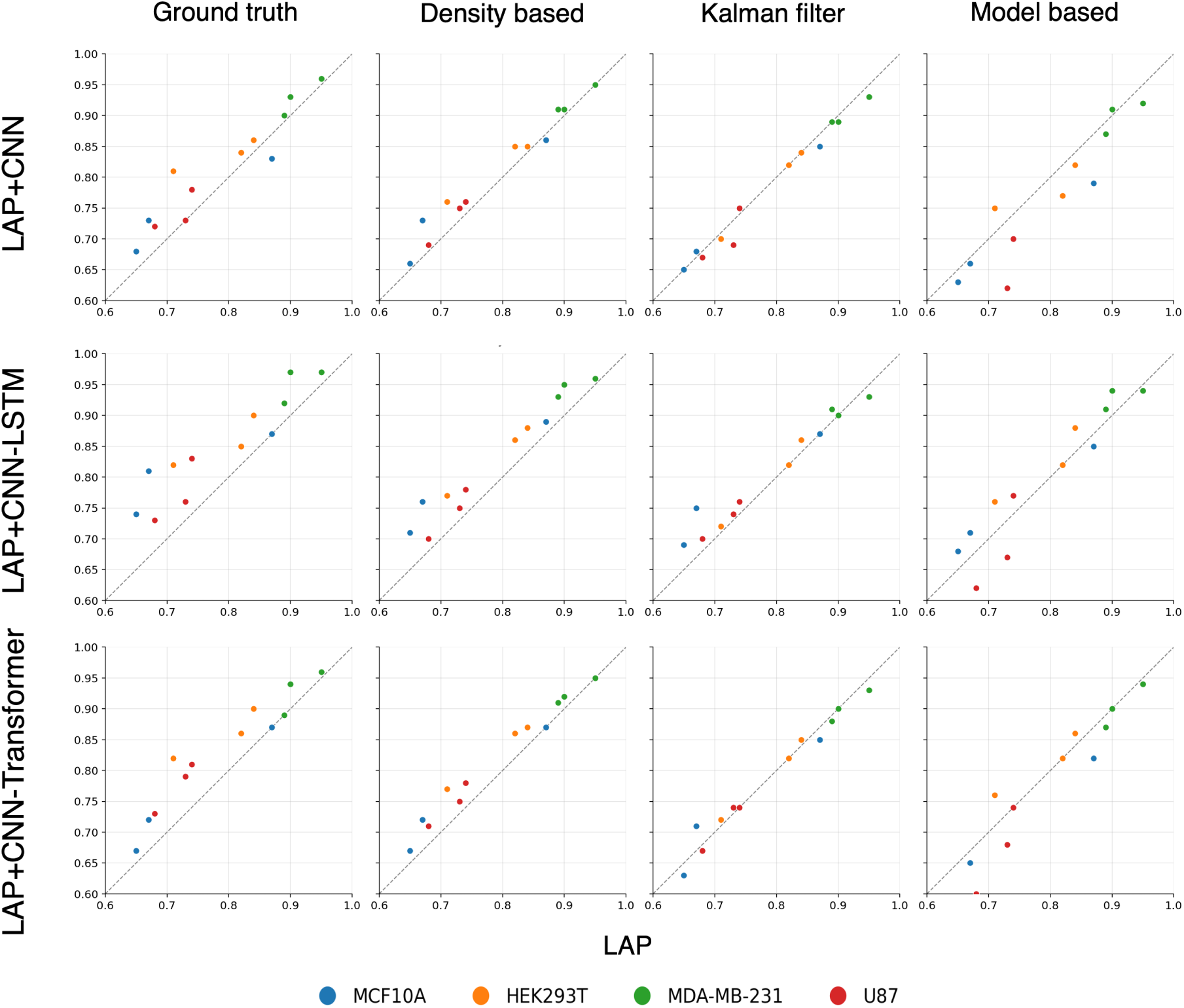
Linking performance of the class-based models at GAP 8. Panels as in Figure S16.

#### S3.2 Continuous-angle

**Figure S18:**
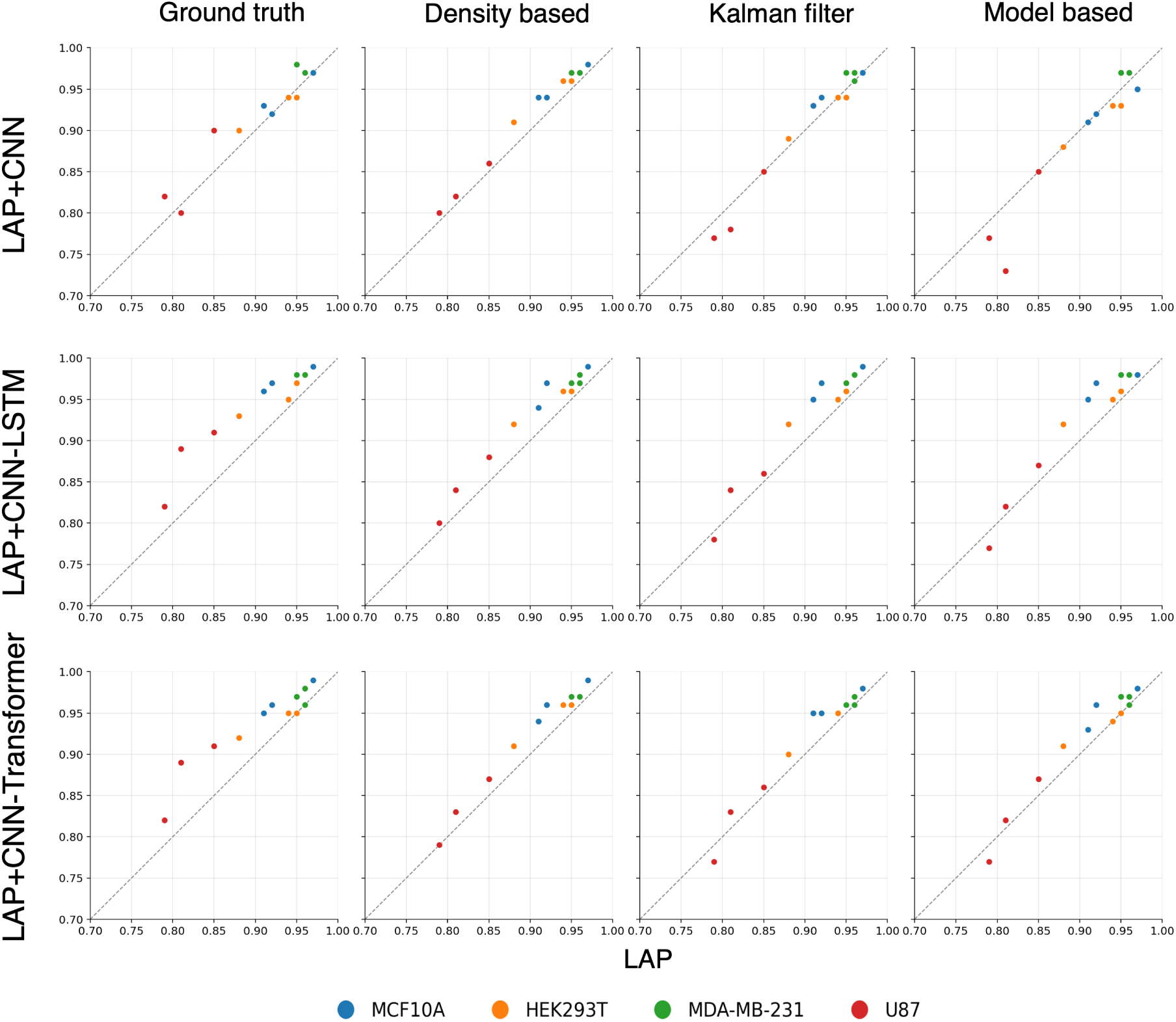
Linking performance of the continuous-angle models at GAP 4. Panels as in Figure S16.

**Figure S19:**
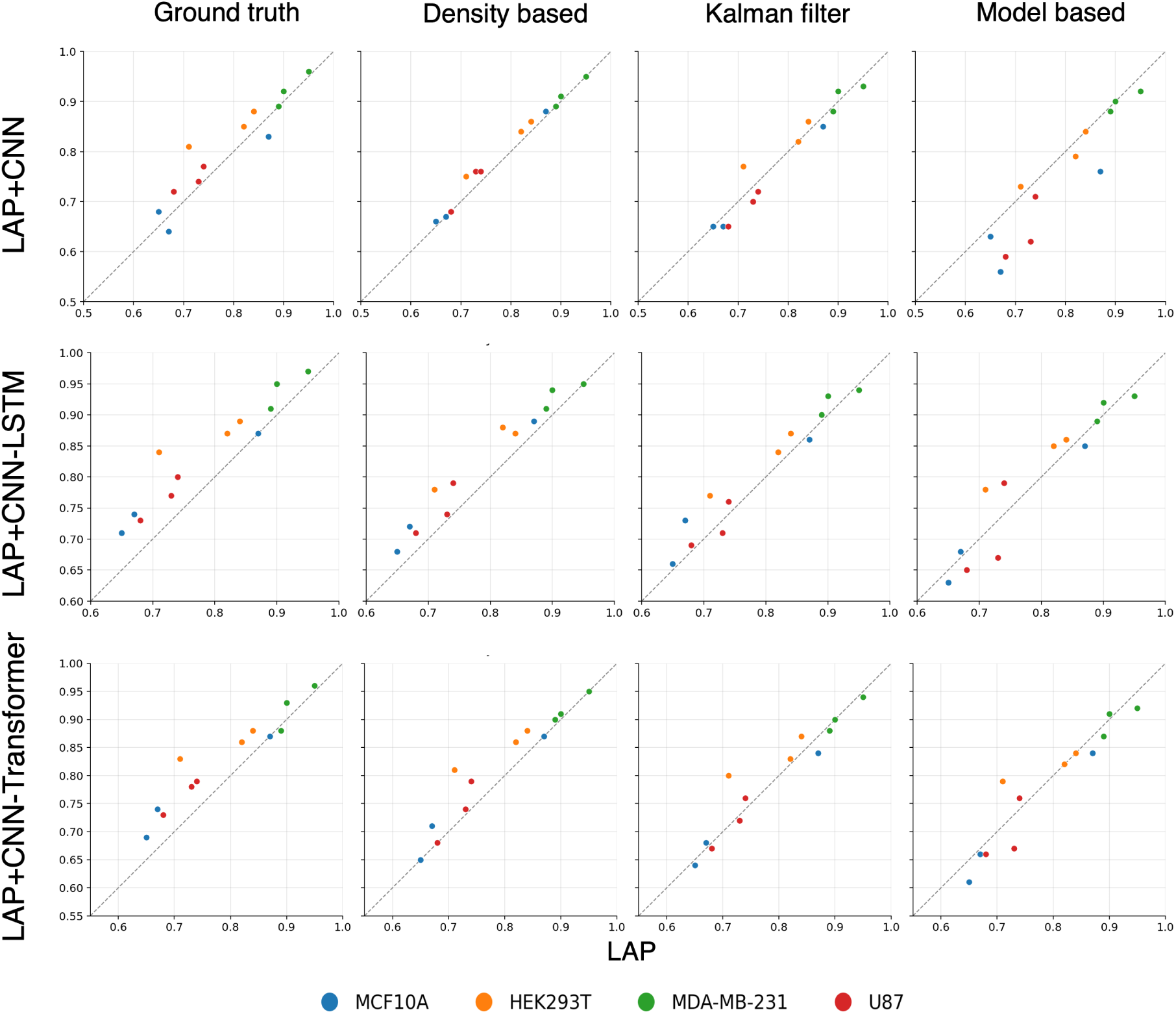
Linking performance of the continuous-angle models at GAP 8. Panels as in Figure S16.

